# BACH2-driven tissue resident memory programs promote HIV-1 persistence

**DOI:** 10.1101/2024.12.16.628794

**Authors:** Yulong Wei, Haocong Katherine Ma, Michelle E. Wong, Emmanouil Papasavvas, Liza Konnikova, Pablo Tebas, Ricardo Morgenstern, Luis J. Montaner, Ya-Chi Ho

## Abstract

Transcription repressor BACH2 redirects short-lived terminally differentiated effector into long-lived memory cells. We postulate that BACH2-mediated long-lived memory programs promote HIV-1 persistence in gut CD4+ T cells. We coupled single-cell DOGMA-seq and TREK-seq to capture chromatin accessibility, transcriptome, surface proteins, T cell receptor, HIV-1 DNA and HIV-1 RNA in 100,744 gut T cells from ten aviremic HIV-1+ individuals and five HIV-1– donors. BACH2 was the leading transcription factor that shaped gut tissue resident memory T cells (TRMs) into long-lived memory with restrained interferon-induced effector function. We found that HIV-1-infected cells were enriched in TRMs (80.8%). HIV-1-infected cells had increased BACH2 transcription factor accessibility, TRM (CD49a, CD69, CD103) and survival (*IL7R*) gene expression, and Th17 polarization (RORC, *CCR6*). In vitro gut CD4+ T cell infection revealed preferential infection and persistence of HIV-1 in CCR6+ TRMs. Overall, we found BACH2-driven TRM program promotes HIV-1 persistence and BACH2 as a new therapeutic target.

## INTRODUCTION

Tissue resident memory T cells (TRMs) protect the gut mucosal barrier against microbial invasion by exerting rapid and localized effector functions while restraining excessive inflammation^1,2^. Despite the absence of antigen stimulation, TRMs persist and remain as long-lived memory cells in tissues^3,4^. TRM T cell development is orchestrated by Hobit and Blimp-1 transcription factors to repress tissue egress genes^5^. Tissue residency is maintained by integrin α1 (*ITGA1*, CD49a)-mediated binding to tissue collagen and laminin and integrin αE (*ITGAE*, CD103)-mediated binding to E-cadherin on the gut epithelium. CD69 expression further prevents egress by inhibiting sphingosine 1-phosphate receptor-1. Although most gastrointestinal infections are transient and self-limiting, HIV-1 establishes major reservoir in gut CD4+ T cells and persists^6–9^. How HIV-1-infected cells survive viral cytopathic effect, evade immune clearance, and persist in the gut remain unknown.

The gut harbors 80–95% of all HIV-1-infected cells^6–9^. Although HIV-1 does not infect gut epithelial cells, HIV-1 replicates in gut CD4+ T cells^7,10–14^, regardless of the route of transmission. Current understanding of mechanisms of HIV-1 persistence is based heavily on CD4+ and CD8+ T cells in the blood^15–18^, where HIV-1-infected cells are believed to be enriched in cytotoxic^15^, activated Th1^19^ (expressing co-receptor CCR5^20^ and CXCR3^21^), Th17^22^, CD161+^23^, central memory (TCM)^24^, and effector memory (TEM)^25^ CD4+ T cells. On the other hand, HIV-1-specific CD8+ T cells are key immune effectors that control HIV-1 infection. HIV-1-specific CD8+ T cells are recruited to the gut^26,27^, may become exhausted during chronic infection^28–30^, and may be rejuvenated after ART^31^. Whether gut HIV-1-specific CD8+ T cells maintain effector functions or become exhausted remains unknown. Given the rarity of HIV-1-infected CD4+ T cells (<0.1%)^15,17,18^ and HIV-1-specific CD8+ T cells, and the requirement of invasive procedures needed to access the gut, understanding mechanisms of HIV-1 persistence and immune evasion in the gut remains challenging.

T cell fate, namely differentiation, effector function, migration, and survival, is governed by transcription factor activity and cytokine cues in the tissue microenvironment. Upon antigen stimulation, gut CD4+ and CD8+ TRMs exert immediate and effective responses to recurrent antigen challenge: Th1 produce interferon γ (IFN-γ) and tumor necrosis factor (TNF) against viral infections, Th17 produce interleukin (IL)-17 and IL-23 against bacterial and fungal infections, Treg produce IL-10 and TGF-β to suppress excessive inflammation and maintain mucosa homeostasis^32,33^, while effector CD8+ T cells secret cytolytic granules containing perforin, granzymes, and granulysin to eliminate infected cells. The majority (>90%) of the terminally differentiated effector T cells die within weeks after antigen stimulation to prevent persistent inflammation^34^. Some cells survive activation-induced cell death and become long-lived memory T cells. We postulate that the dynamic cell fate transition from short-lived terminally differentiated effector to long-lived long-term memory promotes the survival of HIV-1-infected cells.

BACH2 is a key transcription factor that regulates cell fate decision between terminally differentiated effectors and long-lived memory cells^35,36^. BACH2 competes with AP-1 binding to enhancers, attenuates AP-1 proinflammatory activity, restrains effector programs, and generate long-lived memory^36^ and stem-like cells^37^. BACH2 and AP-1 family transcription factors can shape tissue resident memory programs, although BACH2 activity was found to be important in lung and skin but not gut^35^. BACH2 is also one of the most frequent HIV-1 integration sites^38–40^. HIV-1 integration into BACH2 leads to aberrant HIV-1-driven BACH2 expression^41^ and drives clonal expansion of HIV-1-infected cells^39,40^. Knocking in HIV-1 reporter in BACH2 genome induces an immunosuppressive Treg phenotype^42^ and decreased HIV-1 expression^43,44^. Although our study does not have HIV-1 integration site information, we wondered how high BACH2 activity affects the survival and persistence of HIV-1-infected cells, regardless of the HIV-1 integration site.

Given the essential role of transcription factors in tissue residency, effector function, and long-lived memory programs, we aim to identify key transcription factors that shape cellular programs of gut T cells and promote HIV-1 persistence. We hypothesize that BACH2 transcription factor activity shapes gut TRM development from short-lived terminally differentiated effectors to long-lived memory, and that HIV-1 takes advantage of this process and survived in cells having high BACH2 activity. We further postulate that CD8+ TRM program attenuates HIV-1-specific CD8+ T cell effector functions so that they fail to eliminate HIV-1-infected cells. To identify the key transcription factors and the resulting immune programs that shape gut T cells and to decipher mechanisms of HIV-1 persistence in the gut, we coupled single-cell DOGMA-seq^45^ with TREK-seq^46^ to simultaneously capture chromatin accessibility, transcriptome, surface proteins, HIV-1 DNA, HIV-1 RNA, and T cell receptors in the same single cells. Using gene regulatory network (GRN) analysis, we found that BACH2 was the most prominent transcription factor that shaped long-lived memory CD4+ and CD8+ TRM programs and restrained effector T cell programs. By adding TCR repertoire profiling to DOGMA-seq, we identified HIV-1-specific and CMV-specific CD8+ T cells in the gut. We found that HIV-1-specific CD8+ T cells were TRMs with attenuated effector functions. By mapping HIV-1 genome to ATAC-seq and RNA-seq reads, we identified the rare HIV-1-infected cells in the gut. While TRMs accounted for around half (55.1%) of gut CD4+ T cells, the majority (80.8%) of HIV-1-infected cells in the gut were TRMs. BACH2 shaped long-lived memory T cell program and promoted the persistence of HIV-1-infected cells in TRM. Finally, *in vitro* validation using HIV-1-infected primary gut CD4+ T cells identified preferential infection and persistence of HIV-1 in CD4+ TRMs. Our study revealed that BACH2-driven tissue resident program promoted HIV-1 persistence in the gut by attenuating HIV-1-specific gut CD8+ T cell effector function and by maintaining the survival of HIV-1-infected long-lived memory CD4+ T cells.

## RESULTS

### Single-cell DOGMA-seq identified single-cell epigenetic, transcriptional, and protein profiles of gut cell subtypes in human colon

To examine the heterogeneous cell types in the gut, we obtained colon biopsy samples from ten PLWH under ART (plasma viral load <200 copies/ml) and five HIV-1– individuals (HIV–) (**Table S1**) for single-cell DOGMA-seq and TREK-seq to obtain chromatin accessibility landscape (by scATAC-seq), cellular transcriptome (scRNA-seq), 156 surface proteins (by antibody-derived tags (ADT)), and T cell receptor (TCR) sequences within the same single cells. DOGMA-seq profiles RNA from the whole cell and thus avoids the bias of capturing different RNA species in the nuclear versus cytoplasmic RNA in commercially available Multiome methods that requires nuclei isolation. Briefly, a total of 10–14 colon biopsies per participant were processed into single-cell suspensions and cryopreserved into aliquots on the day of biopsy^47^. One aliquot was used for single-cell profiling to examine the proportion of gut immune cells. Another aliquot was enriched for T lymphocytes by magnetic positive selection of CD3+ T cells to examine the T cell programs that promote HIV-1 persistence.

Collagenase treatment is required to isolate immune cells from the lamina propria layer. However, collagenase treatment may degrade surface protein epitopes and reduce surface protein staining^48^. To avoid inaccurate immune profiling because of false negative surface protein staining, we additionally performed DOGMA-seq on peripheral blood mononuclear cells (PBMC) treated with mock, collagenase II, or collagenase IV. We found that collagenase II treatment significantly reduced detection of T cell surface proteins such as CD4, CD45RA, CD183 (encoded by *CXCR3*), CD185 (encoded by *CXCR5*), CD56, and CD119 (encoded by *IFNGR1*) (**Figure S1A, S1B**), whereas collagenase IV treatment significantly reduced CD183 (*CXCR3*) surface protein expression (**Figure S1C**). As collagenase treatment may significantly impact staining of some cell surface proteins, results from protein expression analyses should be cautiously interpreted.

After bioinformatic removal of low-quality cells and doublets, we captured the chromatin accessibility landscape, cellular transcriptome, and surface protein expression in the same single cells, including 17,642 cells from PLWH and 22,422 cells from HIV– individuals in the gut cell aliquot. Combining gut cell aliquot and CD3+ T enriched gut cell aliquot, we captured 43,113 CD4+ T cells from PLWH, 6,092 CD4+ T cells from HIV– individuals, 45,475 CD8+ T cells from PLWH, and 6,064 CD8+ T cells from HIV– individuals. After batch effect removal and weighted nearest neighbor (WNN) integration (by ATAC and RNA profiles), the gut cell phenotypes were visualized by Uniform Manifold Approximation and Projection for Dimension Reduction (UMAP)^49^ (**Figure 1A, Figure S2A**). Surface protein profiles were not considered for WNN integration because collagenase treatment may adversely affect protein expression profiles. We identified 16 gut cell subsets, including CD4+ T cells (16.85%), CD8+ T cells (16.12%), B cells (15.39%), IgA-producing plasma cells (39.10%), IgG-producing plasma cells (19.09%), Innate immune cells (natural killer cells, mast cells, and monocytes) (2.24%), tissue structural cells (epithelial, subepithelial, endothelial, and myofibroblast) (6.39%), and enteric nervous system (myenteric ganglia and enteric glia) (1.57%) (**Figure 1A**, **Table S2**). There was no significant difference in gut cell type proportions between PLWH and HIV– individuals (**Figure 1B**) or between participants (**Figure 1C**).

**Figure 1.**
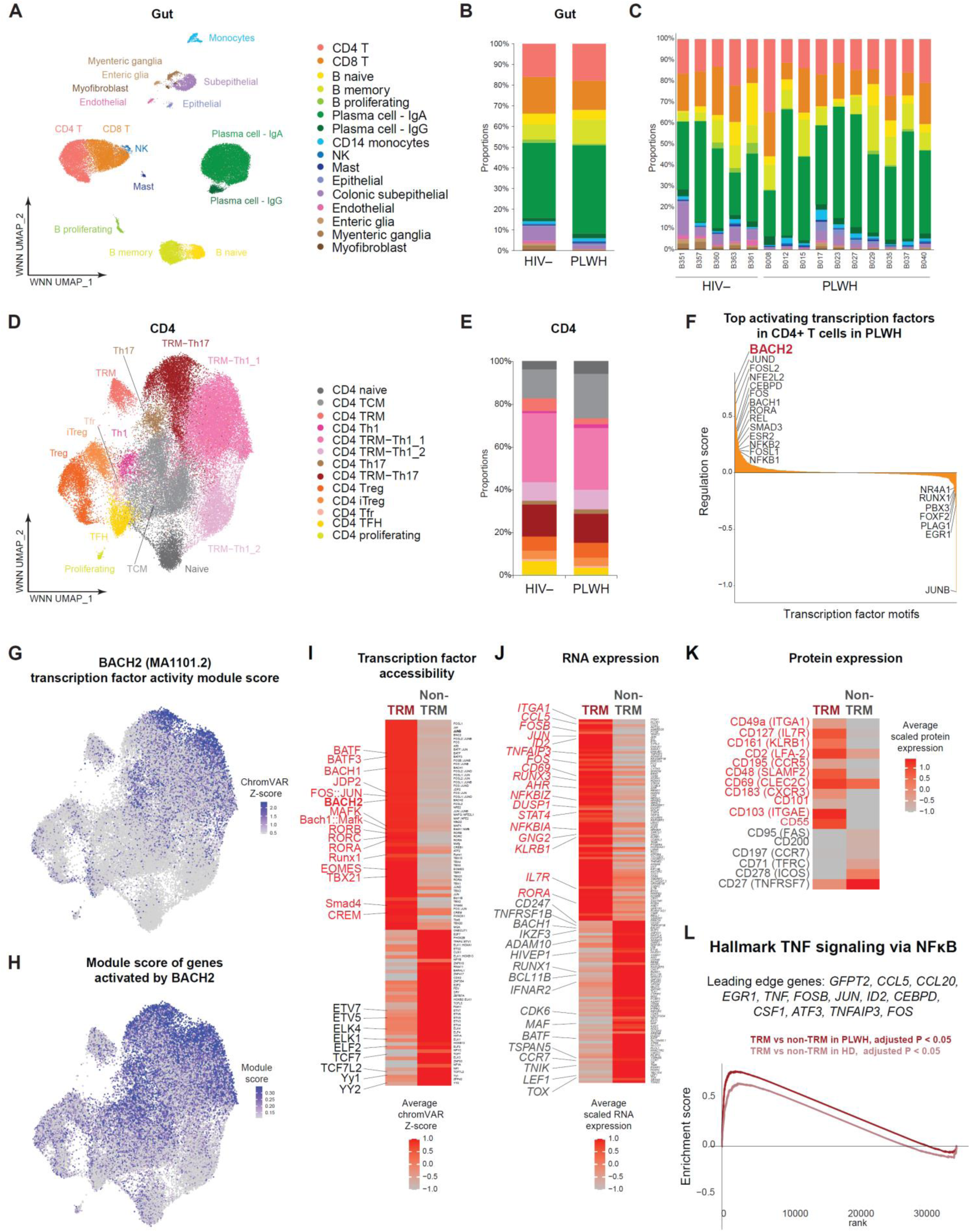
BACH2 drives tissue resident memory CD4+ T cells in human gut. (**A**) WNN UMAP plot of distinct cell subsets in human gut (n = 17,642 in PLWH and 22,422 in HIV– individuals). (**B**) Proportions of gut cell subsets in PLWH and HIV– individuals. (**C**) Proportions of gut cells subsets per participant. (**D**) WNN UMAP plot of distinct CD4+ T cell subsets in human gut (n = 43,113 in PLWH and 6,092 in HIV– individuals), including CD4+ T cells from gut cell aliquots and CD3+ enriched aliquots. (**E**) Proportions of gut CD4+ T cell subsets PLWH and HIV– individuals. (**F**) Mean regulation scores (signed, –log10 scale) across domains of regulatory chromatin (DORC)(n = 639) per transcription factor (TF)(n = 95) for all significant TF-DORC interactions determined in PLWH. Top TF activators (left) and top TF repressors (right) are highlighted. (**G**) WNN UMAP plot of BACH2 transcription factor accessibility in gut CD4+ T cells, measured by chromVAR bias-corrected deviations. (**H**) WNN UMAP plot of per cell RNA module scores in gut CD4+ T cells from PLWH. The module score was built on DORC genes that had upregulated RNA expression in PLWH and were predicted to be activated by BACH2 (BACH2 regulation scores > 0). (**I**) Heatmap indicating differential transcription factors binding motifs accessibility (measured by chromVAR bias-corrected deviations) between CD4+ TRM cells and non-TRM CD4+ cells. FDR-adjusted *P* < 0.05, average Z-score difference > 0.2. (**J**) Heatmap indicating differentially expressed genes (normalized and scaled) between CD4+ TRM cells and non-TRM CD4+ cells. FDR-adjusted *P* < 0.05, min.pct. > 0.25, log2FC > 0.6. (**K**) Heatmap indicating differentially expressed surface proteins (normalized and scaled) between CD4+ TRM cells and non-TRM CD4+ cells. FDR-adjusted *P* < 0.05, min.pct. > 0.15, log2FC > 0.2. The mean expression of all protein features shown were tested to be greater than the mean expression of their specific isotype controls (*Z* > 2; two-sample Z test) in CD4+ T cells. (**L**) To identify enriched pathways in CD4+ TRM T cells, all 36,601 genes were ranked by log2-fold change in normalized gene expression between CD4+ TRM cells and non-TRM CD4+ cells. Significantly enriched TNF signaling pathway was identified using GSEA with leading-edge genes shown. See also Figure S1 – S3.

### BACH2 drives tissue resident memory programs in gut CD4+ T cells

In 43,113 CD4+ cells from PLWH and 6,092 CD4+ T cells from HIV– individuals, we identified 13 CD4+ T cell subsets based on the chromatin accessibility of key transcription factors, RNA expression, and surface protein markers (**Figure S2B – S2D, Table S2**). These CD4+ T cells had high quality profiles in all three modalities with a median of 3,043 unique ATAC fragments, 3,627 ATAC unique molecular identifiers (UMIs), 1,002 genes, 1909 RNA UMIs, 66 surface proteins, and 362 protein UMIs. Overall, 55.09% of CD4+ T cells were tissue resident memory T cells (TRMs), evidenced by high *ITGA1* (integrin α1, an adhesion molecule that binds to collagen) RNA expression, high *CD69* RNA expression, high CD49a (encoded by *ITGA1*) protein expression, and/or high CD103 (integrin αEβ7, E-cadherin-binding molecule encoded by *ITGAE*) protein expression^50,51^. Among CD4+ TRMs, there were four clusters, including two TRM Th1 clusters (TRM Th1_1 had higher levels of *IL12RB2* and *KLRB1*, while TRM Th1_2 had higher levels of *IFNG* and *TNF*), one TRM Th17 cluster, and one TRM cluster. Among CD4+ non-TRMs, there were seven clusters, including naïve, proliferating, central memory (TCM), Th1, Th17, regulatory T cells (Treg), induced Treg (iTreg), follicular regulatory T cells (Tfr), and T follicular helper cells (TFH) (**Figure 1D**). There was no significant difference in CD4+ T cell subset proportions between PLWH and HIV– individuals (**Figure 1E**).

The advantage of having transcription factor activity (by scATAC-seq) and RNA expression (by scRNA-seq) in the same single cells is the ability to unbiasedly identify key transcriptional regulators that governed gene expression by gene regulatory network (GRN) analysis. By GRN analysis, we found that the key transcription factor that regulated gene expression in gut CD4+ T cells was BACH2, followed by AP-1 (JUND, JUN, FOS) (**Figure S3A**). Indeed, BACH2 was a major activator of gene expression in gut CD4+ T cells in PLWH (**Figure 1F**). We found increased accessibility of BACH2 transcription factor motif (JASPAR2022 MA1101.2) (suggesting increased BACH2 transcription factor binding activity) in cells from two TRM populations, TRM-Th17 and TRM Th1_1 (**Figure 1G**). BACH2 accessibility is significantly higher (*P* < 0.05, Wilcoxon rank-sum test) in TRMs in comparison to non-TRMs, in both PLWH and HIV– individuals (**Figure S3B**). Genes predicted to be driven by BACH2 included homing and adhesion molecules integrin α1 (*ITGA1*) and integrin αE (*ITGAE*), cytokine receptors *IL2RB*, *IL4R*, *IL7R*, *IL21R*, *IL23R*, chemokine receptor *CCR7*, and granzyme B inhibitor *SERPINB9* (see **Table S3**). These genes had increased chromatin accessibility to BACH2 binding motif, positive association between *BACH2* expression and target gene’s chromatin accessibility, and positive association between target gene’s chromatin accessibility and RNA expression (see Methods). In addition, these genes had enriched RNA expression in TRM-Th17 and TRM-Th1_1 cells (**Figure 1H**) and significantly higher RNA expression in PLWH versus HIV– individuals and in PLWH TRM vs HIV– individuals TRM (**Figure S3C**) as measured by gene expression module score. Finally, we identified that BACH2-driven genes in the gut were involved in Th1 and Th17 cell differentiation, TCR signaling pathway, and TNF signaling pathway (**Figure S3D**).

### Tissue resident CD4+ T cells do not exhibit effector programs

To identify distinct cellular states of CD4+ TRMs in PLWH, we profiled chromatin accessibility, cellular transcriptome, and surface protein profiles between CD4+ TRMs versus CD4+ non-TRMs in PLWH. Transcription factor activity (as measured by chromatin accessibility of transcription factor binding motifs) was significantly higher in TRMs for transcription factors BACH1 and BACH2, AP-1 family transcription factors (FOS:JUN, JPD2), Th17 master transcription factor RORγt (RORC), Th1 master transcription factor T-bet (TBX21) and EOMES (**Figure 1I**). Significantly upregulated genes in TRMs included AP-1 transcription factors (*FOSB*, *FOS*, and *JUN*), NF-κB inhibitors (*NFKBIA* and *NFKBIZ*), TNF-induced gene *TNFAIP3*, cytokine receptor *IL7R*, Th1 transcription factor *STAT4*, chemokine *CCL5*, and *KLRB1* (**Figure 1J**). Significantly upregulated proteins in TRMs included tissue resident memory markers (CD69, CD49a, and CD103), Th1 markers CXCR3 and CCR5, and Th17 marker CD161^52^ (encoded by *KLRB1*) (**Figure 1K**). Effector T cell programs (involving *GZMA*, *GZMB*, *PRF1*, *ZEB2*, *NKG7*, *IFNG*, and *TNF*) were not upregulated in TRMs. Indeed, genes involved in tissue resident memory (*BACH2, ITGA1*, *CD69*, and *ITGAE*) (**Figure S3E**), Th1 program (*TBX21*, *CXCR3*, *CCR5*, and *CCL5*) (**Figure S3F**), and Th17 program (*RORC*, *CCR6*, *CCL20*, and *KLRB1*) (**Figure S3G**) had increased chromatin accessibility and RNA expression in TRMs. Finally, Gene Set Enrichment Analysis (GSEA) identified significant enriched immune programs in TRMs, such as TNF signaling response (**Figure 1L**) and interferon gamma response (**Figure S2E**). Together, we found that BACH2-driven tissue resident memory CD4+ T programs increased tissue homing, upregulated cytokine receptors and TNF and interferon gamma responses, but not effector programs.

### BACH2 drives tissue resident memory programs in gut CD8+ T cells

After batch effect removal and WNN integration (**Figure S4A**), we identified 11 gut CD8+ T cell clusters in 45,475 gut CD8+ T cells from PLWH and 6,064 gut CD8+ T cells from HIV– individuals (**Figure 2A**) by chromatin accessibility of key transcription factor activity, transcriptome profile, and surface protein expression (**Figure S4b – S4d, Table S2**). Gut CD8+ T cells had a median of 3,230 unique ATAC fragments, 3,884 ATAC UMIs, 987 gene RNA expression, 1,875 RNA UMIs, 67 surface proteins, and 383 UMIs. Overall, 55.98% of CD8+ T cells were TRMs across three clusters (TRM_1, TRM_2, and TRM Tc17) (**Figure 2B**), evidenced by high homing and adhesion molecule expression (integrin αE (*ITGAE*) and integrin α1 (*ITGA1*) gene expression and high CD103 (encoded by *ITGAE*) protein expression) (**Figure S4C, S4D**). In non-TRM CD8+ T cells, we identified seven clusters, including naïve, activated, proliferating, central memory (TCM), effector memory (TEM), γδT, and mucosal-associated invariant T cell (MAIT) CD8+ T cells. In addition, we identified a subset of natural killer T (NKT) cells. There was no significant difference in gut CD8+ T cell subset proportions between PLWH and HIV– individuals (**Figure 2B**).

**Figure 2.**
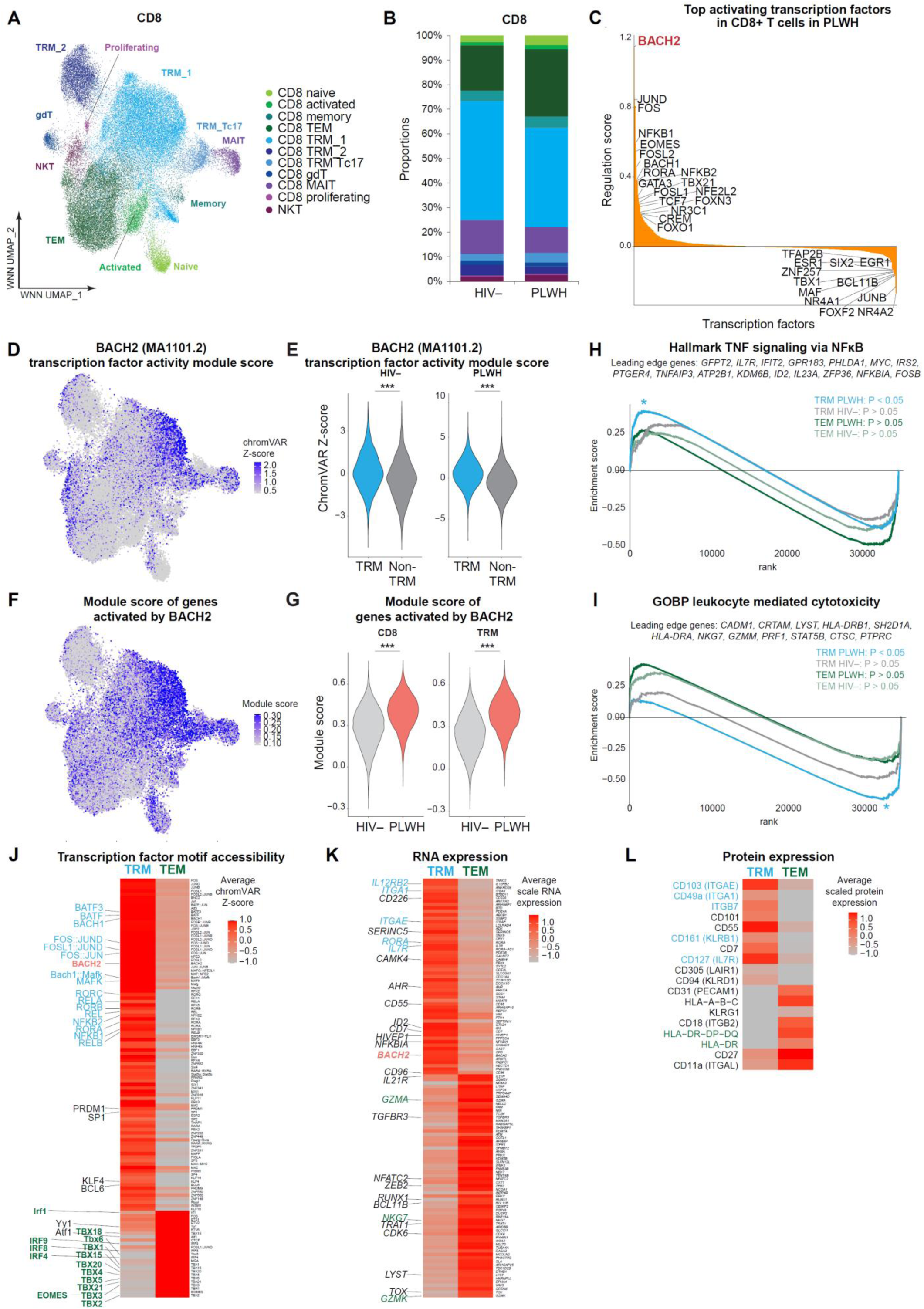
BACH2 drives tissue resident memory program of CD8+ T cells in human gut while interferon regulatory factors (IRF) drive effector programs. (**A**) WNN UMAP plot of distinct CD4+ T cell subsets in human gut (n = 45,475 in PLWH and 6,064 in HIV– individuals), including CD4+ T cells from gut cell aliquots and CD3+ enriched aliquots. (**B**) Proportions of CD8+ T cell subsets in PLWH and HIV– individuals. (**C**) Mean regulation scores (signed, –log10 scale) across DORCs (n = 671) per TF (n = 105) for significant TF-DORC interactions determined in PLWH. (**D**) BACH2 transcription factor accessibility in CD8+ T cells, measured by chromVAR bias-corrected deviations. (**E**) Differences in BACH2 transcription factor accessibility in CD8+ T cells between TRM and non-TRM cells in PLWH and HIV– individuals. (**F**) WNN UMAP plot of per cell module scores. The module score was built on DORC genes that had upregulated RNA expression in PLWH and were predicted to be activated by BACH2 (BACH2 regulation scores > 0) in PLWH CD8+ T cells. (**G**) Differences in module score of DORC genes (upregulated in PLWH and predicted to be activated by BACH2) in CD8+ T cells and CD8+ TRM cells between PLWH and HIV– individuals. (**H,I**) To identify enriched pathway in CD8+ T cell, all 36,601 genes were ranked by log2-fold change in normalized gene expression between CD8+ TRM cell and CD8+ TEM cells in PLWH and in HIV– individuals. (**H**) TNF signaling via NF-κB was enriched in TRM. (**I**) Leukocyte mediated cytotoxicity was enriched in TEM. Significantly enriched pathways were identified using GSEA with leading-edge genes shown. (**J**) Heatmap showing differential transcription factor binding motif accessibility (measured by chromVAR bias-corrected deviations) between CD8+ TRM cells and CD8+ TEM cells. FDR-adjusted *P* < 0.05, average Z-score difference > 0.15. Heatmap indicating differentially expressed genes (normalized and scaled) between CD8+ TRM cells and CD8+ TEM cells (middle). FDR-adjusted p < 0.05, min.pct > 0.25, log2FC > 0.6. (**K**) Heatmap indicating differentially expressed surface proteins (normalized and scaled) between CD8+ TRM cells and CD8+ TEM cells. FDR-adjusted *P* < 0.05, min.pct > 0.15, log2FC > 0.25. (**L**) The mean expression of all protein features shown were tested to be greater than the mean expression of their specific isotype controls (*Z* > 2; two-sample Z test) in CD8+ T cells. ∗ p < 0.05, ∗∗ p < 0.01, ∗∗∗ p < 0.001, Wilcoxon rank-sum test. See also Figure S4.

GRN identified key transcription factors that regulated gut CD8+ T cell programs. We found that BACH2 was also a key regulator (**Figure 2C, Figure S4E**) that shaped gene expression in PLWH gut CD8+ T cells. BACH2 transcription factor activity (measured by binding motif accessibility) was increased in CD8+ TRM_1 (**Figure 2D**), and BACH2 was significantly more accessible (*P* < 0.05, Wilcoxon rank-sum test) in TRMs in comparison to non-TRMs in both PLWH and HIV– individuals (**Figure 2E**). As in CD4+ T cells, genes that were driven by BACH2 included homing and adhesion molecules integrin α1 (*ITGA1*) and integrin αE (*ITGAE*), cytokine receptors *IL2RB*, *IL4R*, *IL7R*, *IL21R*, *IL23R*, chemokine receptor *CCR7*, and granzyme B inhibitor *SERPINB9* (See **Table S3**). These genes were enriched in CD8+ TRM_1 (**Figure 2F**) and significantly upregulated in CD8+ TRMs in PLWH than CD8+ TRMs in HIV– individuals (**Figure 2G**) as measured by gene expression module score. Together, these results suggest that BACH2 is a major driver of tissue resident memory CD8+ T cell program involving tissue homing molecules and cytokine receptor expression.

To examine how BACH2 shaped CD8+ TRMs, we compared the cellular profiles between BACH2-high (high BACH2 binding motif accessibility) CD8+ TRM_1 vs BACH2-low (low BACH2 binding motif accessibility) CD8+ TEM in PLWH. Compared to CD8+ TEM, GSEA identified that CD8+ TRM_1 had significantly enriched TNF signaling responses (**Figure 2H**) and decreased cytotoxicity (**Figure 2I**) in PLWH but not in HIV– individuals. We next profiled chromatin accessibility, cellular transcriptome, and surface protein profiles between CD8+ TRM_1 and CD8+ TEM in PLWH. We found that CD8 TRM_1 had significantly enriched transcription factor activity (as measured by increased binding motif accessibility) for BACH1 and BACH2, AP-1 family transcription factors (FOS:JUN, FOS:JUND, FOSL1:JUND), retinoid acid-related (RAR) orphan receptors RORα (RORA) and RORγt (RORC), NF-κB (RELA, REL, NFKB2, NFKB1, RELB), and follicular T cell lineage defining transcription factor BCL6, while CD8+ TEM had significantly increased transcription factor activity for interferon regulatory factors (IRFs), T-box, and EOMES in CD8+ TEM (**Figure 2J**). Significantly upregulated genes in CD8+ TRM_1 included integrins *ITGA1* and *ITGAE*, cytokine receptors *IL12RB2* and *IL7R*, NF-κB inhibitor *NFKBIA*, whereas significantly upregulated genes in CD8+ TEM included effector molecules *GZMA*, *GZMK*, *ZEB2*, and *NKG7* (**Figure 2K**). Significantly upregulated proteins in CD8+ TRM_1 included markers of tissue resident memory (CD103 and CD49a), cytokine receptor and hallmark of long-lived memory CD127 (encoded by *IL7R*), and CD161 (encoded by *KLRB1*), whereas significantly upregulated proteins in CD8+ TEM included activation markers (HLA-DR, DP, DQ and HLA-A, B, C) and effector T cell marker KLRG1 (**Figure 2L**). Together, our results suggest that BACH2 may regulate TRM programs of both CD4+ T cells and CD8+ T cells in PLWH. TRMs having high BACH2 transcription factor activity were upregulated cytokine receptors for long-lived memory but were downregulated in effector T cell programs.

### BACH2-driven tissue resident memory T cell program maintains long-lived memory cells and restrains terminal effector function

BACH2 is known to drive tissue resident memory^35^, restrain terminal differentiation, and enable generation of long-lived memory CD8+ T cells after murine lymphocytic choriomeningitis virus (LCMV) infection^36^. BACH2 competes with AP-1 transcription factor binding and attenuates T cell activation^36^. However, in the absence of antigen stimulation during ART, TRMs may be natural long-lived memory cells^53^ regardless of BACH2 transcription factor activity. Although BACH2 activity is higher in TRMs than non-TRMs, we want to decipher how BACH2 shaped immune cell phenotype within TRMs. For example, there are two TRM Th1 clusters, both exhibit tissue memory phenotype and Th1 polarization. To determine the role BACH2 played in shaping tissue resident memory T cell in PLWH, we profiled changes in cellular states of TRMs from BACH2-low to BACH2-high CD4+ TRM Th1 (**Figure 3A**) and CD8+ TRM (**Figure 3B**) cells. We want to determine how low versus high BACH2 activity shaped the same immune cell type into two distinct states. To this end, we measured BACH2 transcription factor binding motif accessibility per cell by chromVAR^54^ Z-score and ranked cells from low to high BACH2 activity (as measured by BACH2 binding motif accessibility). Diffusion heatmap was applied to illustrate per-cell feature expression for PLWH cells ranked by BACH2 activity (**Figure 3C – 3F**).

**Figure 3.**
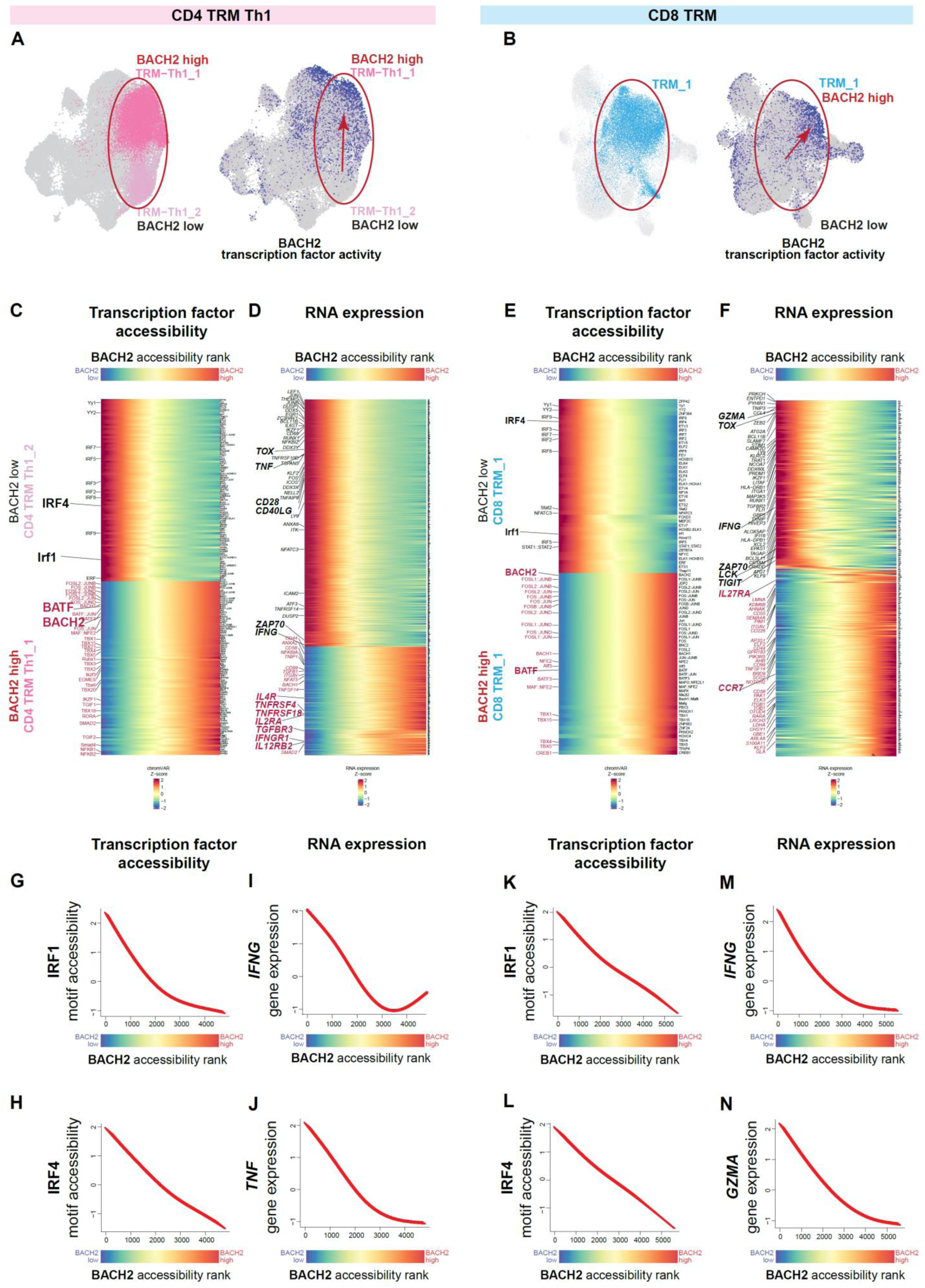
BACH2 restrains terminal effector programs and drives increased long-lived memory cytokine receptor expression in tissue resident memory CD4+ and CD8+ T cells. (**A**) WNN UMAP plot highlighting CD4 TRM Th1_1 and CD4 TRM Th1_2 cells (left) and BACH2 transcription factor accessibility in CD4+ T cells, measured by chromVAR bias-corrected deviations. (**B**) WNN UMAP plot highlighting CD8 TRM_1 cells (left) and BACH2 transcription factor accessibility in CD8+ T cells, measured by chromVAR bias-corrected deviations. (**C – F**) Diffusion heatmap indicating dynamic changes in CD4+ TRM Th1 cells and CD8+ TRM cells ordered from low to high BACH2 accessibility (by chromVAR deviations). Cells were binned into 10 groups by BACH2 accessibility and enriched features were determined in each bin. (**C**) Changes in global chromatin accessibility of transcription factors (measured by chromVAR bias-corrected deviations) from low BACH2 accessible CD4+ TRM Th1 cells (TRM Th1_2) to high BACH2 accessible CD4+ TRM Th1 cells (TRM Th1_1). FDR-adjusted *P* < 0.05, average Z-score difference > 0.3. (**D**) Changes in normalized and scaled gene expression from low BACH2 accessible CD4+ TRM Th1 cells (CD4 TRM Th1_2) to high BACH2 accessible CD4+ TRM Th1 cells (CD4 TRM Th1_1). FDR-adjusted p < 0.05, min.pct. > 0.1, log_2_FC > 0.5. (**E**) Changes in global chromatin accessibility of transcription factors (measured by chromVAR bias-corrected deviations) from low BACH2 accessible CD8+ TRM_1 cells to high BACH2 accessible CD8+ TRM_1 cells. FDR-adjusted *P* < 0.05, average Z-score difference > 0.3. (**F**) Changes in normalized and scaled gene expression from low BACH2 accessible CD8+ TRM_1 cells to high BACH2 accessible CD8+ TRM_1 cells. FDR-adjusted *P* < 0.05, min.pct. > 0.1, log_2_FC> 0.4. (**G – N**) Dynamic feature expression changes in TRM cellular profiles with increase in BACH2 transcription factor accessibility. Accessibility of IRFI (**G**) and IRF4 (**H**) transcription factor binding motifs decreased with increase in BACH2 motif accessibility in CD4+ TRM Th1 cells (TRM Th1_1 and TRM Th1_2 combined). Cytokine *IFNG* (**I**) and *TNF* (**J**) RNA expression decreased with increase in BACH2 motif accessibility in CD4+ TRM Th1 cells (TRM Th1_1 and TRM Th1_2 combined). Accessibility of IRFI (**K**) and IRF4 (**L**) transcription factor binding motifs decreased with increase in BACH2 motif accessibility in CD8+ TRM cells. *IFNG* (**M**) and *GZMA* (**N**) RNA expression decreased with increase in BACH2 motif accessibility in CD8+ TRM cells. See also Figure S5.

We compared the differences in transcription factor activity and gene expression between BACH2-low and BACH2-high CD4+ TRM Th1 (CD4+ TRM Th1_2 and CD4+ TRM Th1_1). The differences in transcription factor activities were that BACH2-low TRM Th1 had higher transcription factor activities for interferon regulatory factors (IRFs), while BACH2-high TRM Th1 had higher transcription factor activities for AP-1, T-box, and NF-κB families (**Figure 3C, 3G, 3H, Figure S5A**). The differences in gene expression were that BACH2-low TRM Th1 had higher levels of effector Th1 cytokines (*TNF* and *IFNG*) and activation molecules (*ICOS*, *CD69*, *CD28*, and *CD40LG*), while BACH2-high TRM Th1 had higher levels of cytokine receptors (*IL2RA*, *IL4R*, *IL12RB2*, *IFNGR1*, *TNFRSF4*, *TNFRSF18*, and *TGFBR3*) (**Figure 3D, 3I, 3J**).

We next compared the differences in transcription factor activity and gene expression between BACH2-low and BACH2-high CD8+ TRM_1. The differences in transcription factor activities were that BACH2-low CD8+ TRMs had higher transcription factor activities for interferon regulatory factors (IRFs) and mediators of IFN signaling STAT1 and STAT2, while BACH2-high CD8+ TRMs had higher transcription factor activities for AP-1 and T-box families (**Figure 3E, 3K, 3L, Figure S5B**). The differences in gene expression were that BACH2-low CD8+ TRMs had higher levels of effector molecules (*GZMA*, *ZEB2*, *IFNG*, *KLRC2*, and *CCL4*) while BACH2-high CD8+ TRMs had higher levels of cytokine receptor *IL27RA*, chemokine receptor *CCR7*, adhesion molecules (*ITGAV*, *ITGB1*) **(Figure 3F, 3M, 3N**).

Overall, within CD4+ and CD8+ TRMs, we found that in BACH2-low cells, interferon responses drove TRMs into effector cells, while in BACH2-high cells, BACH2 drove TRMs into tissue residence (adhesion molecules) and long-lived memory cells (upregulating receptors that promote survival while downregulating effector molecules). Our results suggest that BACH2 transcription factor activity maintains long-lived memory and tissue retention and restrains terminal effector function and activation-induced cell death.

### TNF and IFN-γ effector cytokines from BACH2-low TRM senders contribute to TNF and IFN-γ responses in BACH2-high, long-lived memory, TRM receivers

Our results showed that in PLWH, BACH2-low effector TRM exhibited hallmarks of reactivated TRMs and expressed *IFNG* and *TNF* cytokines^1,2^, whereas BACH2-high, long-lived, TRMs expressed cytokine receptors and TNF- and IFN-γ-induced genes. We speculated that the cytokine ligand-receptor interactions between sender and receiver cells, i.e. TNF and IFN-γ effector cytokine secretion from effector cells to downstream cytokine receiving long-lived memory cells, promoted the persistence of long-lived memory cells. We performed intercellular communication analysis between all gut CD4+ and CD8+ T cells for an unbiased discovery of potential ligand – receptor interaction pairs. We merged CD4+ T cell and CD8+ T cell datasets, performed batch-effect correction and WNN integration, and visualized these cells on UMAP (**Figure 4A**). Using Scriabin^55^, we uncovered two statistically significant ligand – receptor interaction programs (modules of ligand – receptor pairs co-expressed by the same ligand-expressing ‘sender’ cells and receptor-expressing ‘receiver’ cells), Interaction Program 2 (IP-2) (**Figure S5C**) and Interaction Program 12 (IP-12) (**Figure S5D**), that were significantly enriched in PLWH in comparison to HIV– individuals. Together, IP-2 and IP-12 identified significant intercellular communication between ligand-expressing activated CD8+ T cells and TRM Th1_2 cells (BACH2-low) and receptor-expressing TRM Th1_1 and TRM Th17 cells (BACH2-high) (**Figure 4B, Figure S5E, S5F**). Among significantly enriched ligand – receptor pairs in PLWH identified from IP-2 (**Figure S5E**) and IP-12 (**Figure S5F**) were *TNF* ligand expression from activated CD8+ T and TRM Th1-2 sender cells and *TNFRSF1A* and *TNFRSF1B* receptor expression from TRM Th1_1 and TRM Th17 receiver cells, and *IFNG* ligand expression from the same sender cells to *IFNGR1* and *IFNGR2* receptor expression from the same receiver cells. Indeed, activated CD8+ T cells and TRM Th1-2 cells highly expressed *IFNG* and *TNF* while TRM Th1_1 and TRM Th17 cells highly expressed *IFNGR1*, *IFNGR2*, and *TNFRSF1B* (**Figure 4C**).

**Figure 4.**
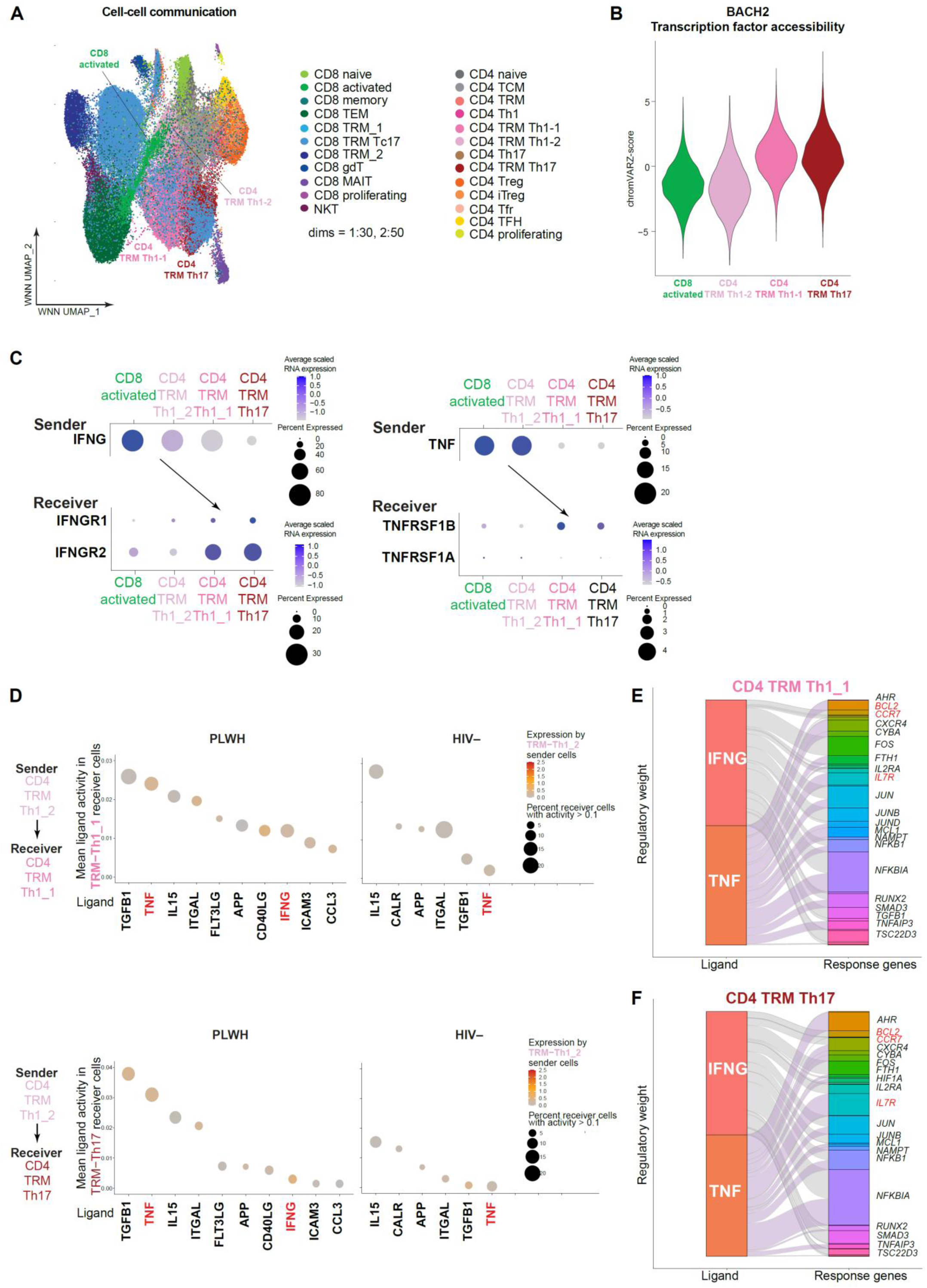
TNF and IFN-γ communication from BACH2-low ligand-expressing sender cells to BACH2-high receptor-expressing receiver CD4+ TRM. (**A**) WNN UMAP plot of distinct CD4+ and CD8+ T cell subsets in human gut (n = 43,113 and 45,475 in PLWH, 6,092 and 6,064 in HIV– individuals, respectively). (**B**) BACH2 transcription factor accessibility (measured by chromVAR bias-corrected deviations) in CD8 activated, CD4 TRM Th1_2, CD4 TRM Th1_1, and CD4 TRM Th17 cells. (**C**) Scriabin interaction program discovery identified *IFNG* ligand to *IFNGR1* and *IFNGR2* receptor interactions (left) and *TNF* ligand to *TNFRSF1B* and *TNFRSF1A* receptor interactions (right) between sender (ligand expressing) CD8 activated and CD4 TRM Th1_2 cells and receiver (receptor expressing) CD4 TRM Th1_1 and CD4 TRM Th17 cells. Dot plots showed average normalized and scaled RNA expression of ligand and receptor genes. (**D**) Dot plot depicting NicheNet-predicted ligand activity in receiver TRM-Th1_1 (top) and TRM Th17 (bottom) in PLWH and HIV– individuals. Dot size depict percentage of receiver cells having ligand activity weight of > 0.1. Color scales represent ligand expression from TRM-Th1_2 sender cells (measured by RNA expression fold change in TRM-Th1_2 relative to other cell clusters). (**E**) Alluvial plot depicting target genes predicted to be upregulated in receiver TRM Th1_1 cells in response to *IFNG* and *TNF*. (**F**) Alluvial plot depicting target genes predicted to be upregulated in receiver TRM Th17 cells in response to *IFNG* and *TNF*. See also Figure S5.

To determine whether *TNF* and *IFNG* ligands shaped receptor-expressing TRM Th1_1 and TRM Th17 cells, we applied NicheNet^56^ and identified increased cellular responses to TNF and IFN-γ signaling (by ligand activity that measured RNA expression of target gene due to ligand – receptor interactions) in both TRM Th1_1 and TRM Th17 receiver cells (**Figure 4D**). In particular, mean ligand activity scores of both *TNF* and *IFNG* in TRM Th1_1 and TRM Th17 receiver cells were higher in PLWH in comparison to HIV– individuals, suggesting that TNF and IFN-γ signaling were increased in TRM Th1_1 and TRM Th17 cells in PLWH. In addition, Scriabin-predicted senders of *TNF* and *IFNG* ligands, activated CD8 (**Figure S5G, S5H**) and TRM Th1_2 (**Figure 4D**), were cells that had the highest *TNF* and *IFNG* gene expression (as shown by color scale representing log2-fold change in RNA expression) among T cells. Finally, NicheNet-predicted target genes that were upregulated as a result of *IFNG* and *TNF* signaling included AP-1 genes *FOS* and *JUN*, NF-κB genes *NFKB*, *NFKBIA*, cytokine and chemokine receptors *IL2RA*, *IL7R*, *CCR7*, survival gene *BCL2* and TNF-induced gene *TNFAIP3* in both receivers TRM-Th1_1 (**Figure 4E**) and TRM Th17 (**Figure 4F**). These results suggest that BACH2-low TRMs (namely TRM Th1_2) maintained effector function and contributed to increased TNF signaling and IFN-γ responses in BACH2-high, long-lived TRMs (namely TRM Th1_1 and TRM Th17). Because of high *TNF* and *IFNG* ligand activities, BACH2-high TRM Th1_1 and TRM Th17 cells had increased RNA expression for cytokine and chemokine receptors (*IL2RA*, *IL7R*, *CCR7*) and survival (*BCL2*).

### Adding TCR profiling to DOGMA-seq identified HIV-1 antigen specific CD8+ T cells in TRM

To identify HIV-1-specific CD8+ T cells in the gut, we used TREK-seq^46^ to identify T cell receptor (TCR) sequences using an aliquot of DOGMA-seq cDNA library. This allowed us to identify TCR sequences and respective ATAC-seq, RNA-seq, and protein profiles within the same single cells. We identified HIV-1-specific versus CMV-specific CD8+ T cells by mapping TCRβ CDR3 junction amino acid sequences against the McPAS-TCR database^57^ of TCRs of known HIV-1 and CMV antigen specificity using three stringent criteria^58^. We captured TCRβ sequences in 10,868 out of 45,475 (23.9%) CD8+ T cells in PLWH and 875 out of 6,064 (14.4%) CD8+ T cells in HIV– individuals. Out of 10,868 TCR-mapped CD8+ T cells in PLWH, we identified 56 HIV-1-specific CD8+ T cells and 429 CMV-specific CD8+ T cells (**Figure S6A**, **Table S2, Table S4**). The majority of HIV-1-specific CD8+ T cells were TRMs (67.86%). In particular, HIV-1-specific CD8+ T cells were enriched in long-lived TRMs (50.00% of HIV-1-specific CD8+ T cells were CD8+ TRM_1, compared to 38.69% for CMV-specific CD8+ T cells and 42.95% for other antigen-specific CD8+ T cells in CD8+ TRM_1) (**Figure S6B**, **Table S2**).

### HIV-1-specific CD8+ T cells have decreased effector molecule expression than CMV-specific CD8+ T cells in the gut

CMV-specific CD8+ T cells were known to retain effector function in HIV-1 infection, while HIV-1-specific CD8+ T cells were thought to be exhausted during chronic infection^59^. Comparing the immune profiles of HIV-1-specific CD8+ T cells versus CMV-specific CD8+ T cells, we found that HIV-1-specific CD8+ T cells had higher levels of TRM markers *ITGAE* and *ITGA1*, cytokine receptor *IL12RB2*, and *KLRB1* (**Figure S6C**). Comparing HIV-1-specific CD8+ T cells versus other antigen specific CD8+ T cells, HIV-1-specific CD8+ T cells still had higher levels of TRM markers *ITGAE* and *ITGA1* expression (**Figure S6D**). Comparing surface protein expression profiles, we found that HIV-1-specific cells had lower level of CD183 (CXCR3) than CMV-specific cells (**Figure S6E**) and other antigen-specific cells (**Figure S6F**).

Because HIV-1-specific CD8+ T cells were enriched in BACH2-high CD8+ TRM_1, we postulated that HIV-1-specific CD8+ T cells have reduced effector responses than CMV-specific CD8+ T cells^59^. At the RNA expression level, we found that HIV-1-specific CD8+ T cells had higher levels of tissue resident *ITGA1* and *ITGAE* expression but lower levels of *GZMA*, *GZMK*, *GZMH*, *ZEB2*, *NKG7*, and *CCL5* effector gene expression than CMV-specific CD8+ T cells and other antigen-specific CD8+ T cells (**Figure S6G**). Of note, granzyme A, as opposed to granzyme B, is key to protective immunity in the gut during *Salmonella* infection, independent of perforin^60^. At the protein expression level, HIV-1-specific CD8+ T cells had higher level of homing marker CD49a (*ITGA1*) but lower levels of activation and immune checkpoint molecules CD150 (SLAMF1), CD279 (PD-1), CD152 (CTLA-4), CD223 (LAG-3), CD39 (*ENTPD1*), and TIGIT in comparison to CMV-specific and other antigen-specific CD8+ T cells (**Figure S6H**). Of note, these trends shown in dot plots should be cautiously interpreted as the number of HIV-1-specific cells captured (n = 56) were too few for robust statistical tests. Overall, our results suggest that HIV-1-specific CD8+ T cells are recruited to the gut TRM. They not exhausted but have reduced T cell effector gene expression than CMV-specific CD8+ T cells.

### HIV-1-specific CD8+ T cells are less proliferative than CMV-specific CD8+ T cells

Upon antigen stimulation, naïve CD4+ T cells and CD8+ T cells are activated and proliferate into many cells having the same TCR sequence, called a T cell clone. When the same antigen stimulation continues or recurs, the corresponding T cell clone proliferate. T cell clones were identified from TCR-mapped cells based on their shared TCRβ CDR3 junction sequences (see Methods), as cells having identical TCRs presumably originate from the same T cells that proliferated into a T cell clone in response to the same antigen stimulation. To determine whether continuous HIV-1 antigen stimulation occurred in PLWH under ART, we found that 25 out of 56 (44.64%) HIV-1-specific CD8+ T cells were in clones, indicating proliferation *in vivo*. In comparison, 259 out of 429 (60.37%) of CMV-specific CD8+ T cells in PLWH were in clones (**Table S2**). Of note, HIV-1-specific clones were detected in only four out of ten PLWH participants (015, 029, 035, 040) with smaller clone size than CMV-specific CD8+ T cells (**Figure S6I**, see **Table S3**), suggesting low antigen stimulation or low proliferation capacity of HIV-specific CD8+ T cells than CMV-specific CD8+ T cells.

### CD4+ T cell clones are less proliferative than CD8+ T cell clones

We wanted to compare the proliferation capacity of CD4+ T cells versus CD8+ T cells. We mapped TCR sequences and measured CD4+ T cell clone size. We identified TCRβ sequences in 13,012 out of 43,113 (30.18%) CD4+ T cells in PLWH, and in 1,104 out of 6,092 (18.12%) CD4+ T cells in HIV– individuals. Unique participant-specific clones were determined by shared TCRβ CDR3 junction sequences for CD4+ T and CD8+ T cells. Comparing clone size (normalized in counts per million cells) between CD4+ T cells and CD8+ T cells in PLWH, we found that CD8+ T cells were more clonally expanded and harbored the largest clones in comparison to CD4+ T cells (**Figure 5A**).

**Figure 5.**
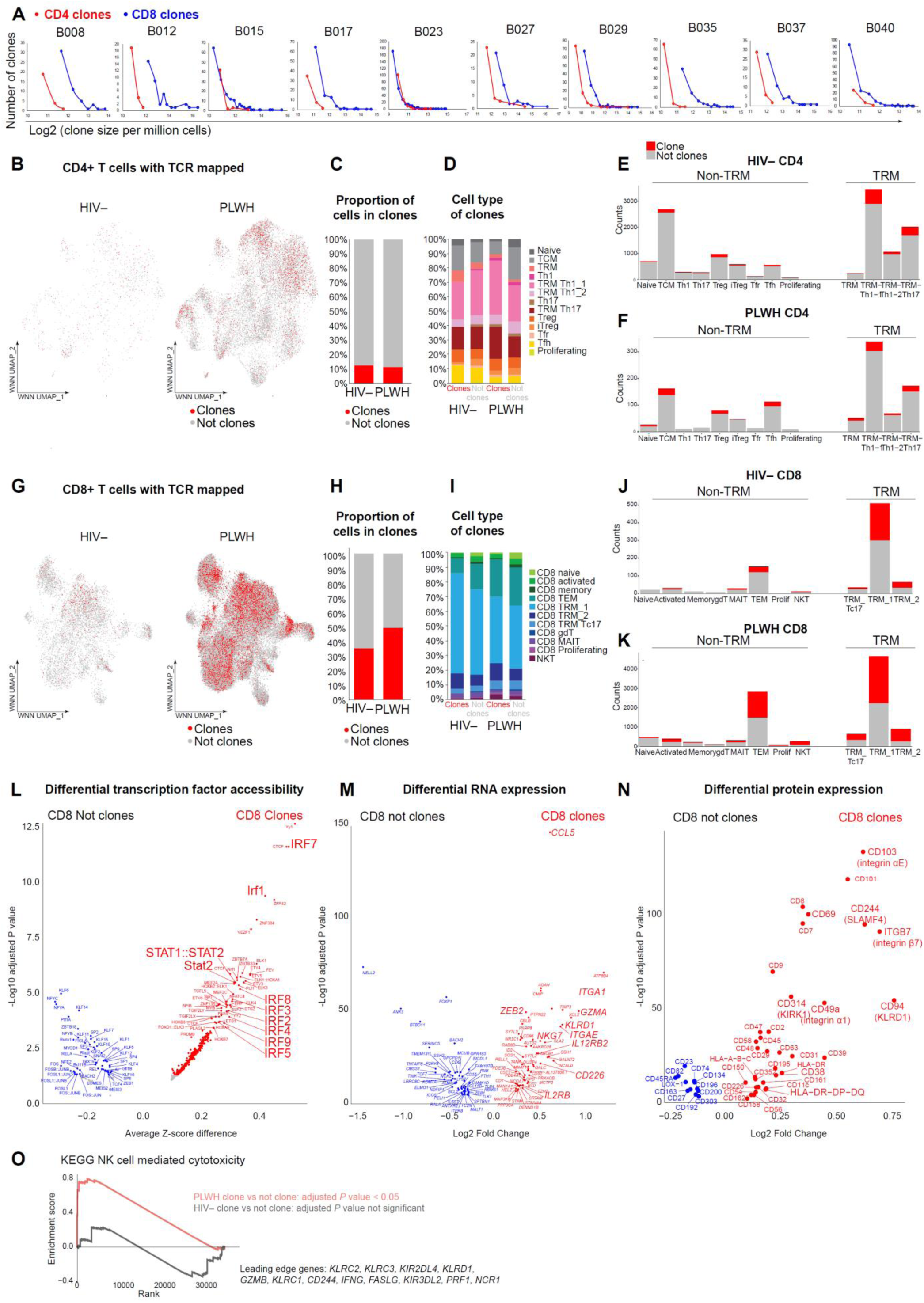
IRF drives clonal expansion of effector CD8+ T cells in the gut. (**A**) Distribution of log-transformed T cell clone size per 1,000,000 cells for all unique T cell clones identified per PLWH participant. **(B)** WNN UMAP plot showing CD4+ T cell clones in PLWH (1485/13,012) and in HIV– individuals (875/1,104). **(C)** Proportions of CD4+ T cell in clones. (**D**) Cell subset proportions of CD4+ T cell clones and non-clones. (**E,F**) Proportions of CD4+ T cell in clones in each subset in HIV– individuals gut (**E**) and PLWH gut (**F**). (**G**) WNN UMAP plot showing CD8+ T cell clones PLWH (5,510/10,868) and in HIV– individuals (308/875). (**H**) Proportions of CD8+ T cell in clones. (**I**) Cell subset proportions of CD4+ T cell clones and non-clones. (**J,K**) Proportions of CD8+ T cell in clones in each subset in HIV– individuals gut (**J**) and PLWH gut (**K**). (**L**) Differential transcription factors binding motif accessibility (measured by chromVAR bias-corrected deviations) between CD8+ T cell clones and non-clones. FDR-adjusted *P* < 0.05, average Z-score difference > 0.1. (**M**) Differentially expressed genes (normalized and scaled) between CD8+ T cell clones and non-clones. FDR-adjusted *P* < 0.05, min.pct > 0.25, log2FC > 0.25. (**N**) Differentially expressed surface proteins (normalized and scaled) between CD8+ T cell clones and CD8+ T cells that were not clonal. FDR-adjusted *P* < 0.05, min.pct > 0.1, log2FC > 0.25. The mean expression of all protein features shown were tested to be greater than the mean expression of their specific isotype controls (*Z* > 2; two-sample Z test) in CD8+ T cells. (**O**) To identify enriched pathways in CD8+ T cell clones, all 36,601 genes were ranked by log2-fold change in normalized gene expression between CD8+ T cell clones and CD8+ cells that were not clonal. Significantly enriched cytotoxicity pathway was identified using GSEA with leading-edge genes shown. All comparisons were made for CD8+ T cells whose CDR3β junction sequences were captured. See also Figure S6 – S8.

### BACH2-high TRM Th1 and TRM Th17 are the most clonally expanded CD4+ T cells in the gut

We next compared CD4+ T cell clone size in PLWH versus HIV– individuals. We identified 1,485 out of 13,012 (11.41%) TCR-mapped CD4+ T cells in clones in PLWH and 138 out of 1,104 (13.6%) TCR-mapped cells in clones in HIV– individuals (**Figure 5B**, **Table S2, Table S4**). There was no significant difference in cell type composition of clones in PLWH versus HIV– individuals (**Figure 5C**). In PLWH, most clones were identified from TRM Th1_1 (37.58%) and TRM Th17 (22.36%), TCM (8.89%), and Treg (8.22%). In HIV– individuals, most clones were identified from TRM Th1_1 (26.09%), TRM Th17 (15.94%), TCM (17.39%), TFH (12.32%), and Treg (8.70%) (**Figure 5D**, **Table S2**). However, there was no clonal proliferation enrichment for any cell subsets (comparable proportions of clones to cells in each cluster) in PLWH and HIV– individuals (**Figure 5E, 5F**).

### TEM and BACH2-high TRM are the most clonally expanded CD8+ T cells in the gut

In CD8+ T cells, we identified 5,510 out of 10,868 (50.70%) TCR-mapped CD8+ T cells in clones in PLWH, and 308 out of 875 (35.2%) TCR-mapped CD8+ T cells in clones in HIV– individuals (**Figure 5G**, **Table S2, Table S4**). Although TCR-mapped HIV– individual cells were limited in cell number, higher proportions of cells were found as CD8+ T cell clones in PLWH (50.70%) than HIV– individuals (35.2%) (**Figure 5H**). By proportions, clonally expanded CD8+ T cells in HIV– individuals were enriched in BACH2-high TRM_1 (68.18% in HIV– individuals vs 45.28% in PLWH), while clonally expanded CD8+ T cells in PLWH were enriched in TEM (25.27% in PLWH vs 10.06% in HIV– individuals) (**Figure 5I**). In both PLWH and HIV– individuals, most clones were identified from TRM_1 (45.28% and 68.18%, respectively), TRM_2 (11.91% and 10.38%, respectively), and TEM (25.27% and 10.06%, respectively) (**Figure 5J, 5K**).

### Clonally expanded gut CD8+ T cells are effectors driven by interferon responses

To identify the immune programs driving the clonal proliferation of gut CD8+ T cells, we identified chromatin accessibility, cellular transcriptome, and cell surface protein profiles of CD8+ T cell clones. We compared CD8+ T cell clones to cells that were not clonal (TCR-mapped cells that were not identified as clones) in PLWH. At the level of transcription regulation, we found that interferon regulatory factors (IRF) and mediators of IFN signaling STAT1 and STAT2 were the prominent transcription factors that were significantly more accessible in CD8+ T cell clones (**Figure 5L**). At the level of RNA expression, we found that effector genes (*GZMA*, *CCL5*, *ZEB2*, *KLRD1*, *NKG7*), cytokine receptors (*CD226* (a TNF receptor encoded by *TNFRSF12A*), *IL2RB*, and *IL12RB2*), and tissue resident adhesion molecules (*ITGAE* and *ITGA1*) were significantly higher in CD8+ T cell clones (**Figure 5M**). At the level of surface protein expression, effector markers [CD56 (NCAM1), CD58 (LFA3), CD94 (KLRD1), CD158 (KIR), CD226, and CD314 (KIRK1)] and T cell activation markers [CD38, CD244 (SLAMF4), CD150 (SLAMF1), HLA-DR, DP, DQ, and HLA-A, B, C], and tissue resident adhesion molecules [CD103 (ITGAE), CD49a (ITGA1), and CD69] were significantly higher in CD8+ T cell clones (**Figure 5N**).

We next focused on gut CD8+ TRM_1 to mitigate noise in cellular profiles introduced by cell type bias of CD8+ T cell clones. We repeated differential expression analyses for CD8+ T cell clones versus CD8+ T cells that were not clonal within TRM_1 cluster in PLWH. We found that clonally proliferated CD8+ T cells within TRM_1 had increased accessibility for IRF7 (**Figure S7A**), increased RNA expression of effector (*CCL5*, *GZMA*, *ZEB2*, *NKG7*) and tissue resident adhesion molecules (*ITGAE* and *ITGA1*) (**Figure S7B**), and increased protein expression of effector (CD56), activation [CD314 (NKG2D), CD244 (SLAMF), CD150 (SLAM), HLA-DR], and tissue resident adhesion molecules [CD49a (ITGA1), CD69, and CD103 (ITGAE)] (**Figure S7C**). Overall, IRF-driven effector program drives the proliferation of CD8+ T cells.

We examined the cellular profiles of CD8+ T cell clones between PLWH and HIV– individuals. We found that CD8+ T cell clones in PLWH had increased accessibility for IRF transcription factors **(Figure S7D**), increased RNA expression of effector genes (*CCL5*, *GZMA*, and *NKG7*) (**Figure S7E**), and increased protein expression of T cell activation makers (CD244 (SLAMF), CD150 (SLAM), HLA-DR, DP, DQ, HLA-A, B, C), and tissue resident adhesion molecules (CD69, CD103) (**Figure S7F**). Indeed, effector genes such as *CCL5* and *GZMA* had significantly increased chromatin accessibility at gene locus and increased RNA expression in CD8+ T cell clones (**Figure S7G**), whereas hallmark genes of long-term memory cells *BACH2* and *CCR7* had significantly decreased chromatin accessibility at gene locus and decreased RNA expression in CD8+ T cell clones (**Figure S7H**). Using Gene Set Enrichment Analysis (GSEA), we found that CD8+ T cell clones in PLWH but not in HIV– individuals had significant enrichment in T cell cytotoxicity (including effector T cell leading genes *KLRC1*, *KLRC2*, *KLRC3*, *KIR2DL1*, *KIR2DL4*, *KIR3DL2*, *KLRD1*, *GZMB*, and *PRF1*) (**Figure 5O**). Furthermore, gene ontology analysis showed that CD8+ T cell clones had significant enrichment of T cell proliferation, interferon gamma production, and killer cell mediated immunity versus cells that are not clonal in PLWH (**Figure S7I**). Lastly, CD8+ T cell clones in PLWH had enriched T cell activation and killer cell mediated immunity versus CD8+ T cell clones in HIV– individuals (**Figure S7J**). Together, we found that CD8+ T cell clones had increased effector responses compared to CD8+ T cells that were not clonal in PLWH and compared to CD8+ T cell clones in HIV– individuals.

### Clonally expanded gut CD4+ T cells exhibit TRM Th1 and Th17 phenotypes

Comparing CD4+ T cell clones to cells that are not clonal in PLWH, we determined that the cellular profiles of CD4+ T cell clones closely resembled that of TRM Th1_1 and TRM Th17 cells. At the transcription regulation level, clonally expanded gut CD4+ T cells had enriched accessibility for AP-1 transcription factors, Th17 master transcriptional regulator RORC, and Th1 master transcriptional regulator TBX21 and EOMES (**Figure S8A**). At the RNA expression level, clonally expanded gut CD4+ T cells had increased levels of *CCL5*, *IL12RB2*, *RORA*, *TNFAIP3*, *KLRB1*, *NFKBIA*, and *SERPINB9* gene expression (**Figure S8B**). At the protein expression level, clonally expanded gut CD4+ T cells had increased levels of tissue resident memory [CD103 (ITGAE) and CD49a (ITGA1)], Th1 [CD195 (CCR5)], and CD161 protein expression (**Figure S8C**). Nonetheless, this finding is not surprising since 59.93% of all CD4+ T cell clones in PLWH were identified in these TRM Th1_1 and TRM Th17 (**Figure 5D**, **Table S2**). To mitigate cellular profile noise introduced by cell type bias of CD4+ T cell clones, we compared CD4+ TRM clones to CD4+ TRMs that were not clonal in PLWH. We still identified enriched accessibility for AP-1 and NF-κB (REL, RELA) transcription factors (**Figure S8D**), upregulated *CCL5*, *ITGA1*, *KLRB1*, and *IFNGR1* gene expression (**Figure S8E**), and CD69, CD103 (ITGAE), CD161 (KLRB1), CD195 (CCR5) protein expression (**Figure S8F**) in CD4+ TRM clones.

Lastly, we compared the cellular profiles of CD4+ T cell clones between PLWH and HIV– individuals. We found that gut CD4+ T cell clones in PLWH had increased accessibility for AP-1, IRF3, and RORC transcription factors (**Figure S8G**), upregulated *CCL5*, *HLA-A*, *HLA-B*, *HLA-C*, and *CD44* gene expression (**Figure S8H**), and upregulated CD69, CD44, CD183 (CXCR3), CD150 (SLAM), CD161 (KLRB1), CD195 (CCR5), and activation (HLA-A, B, C, and HLA-DR, DP, DQ) protein expression (**Figure S8I**). Together, we identified that CD4+ T cell clones exhibit long-lived memory phenotypes that highly resembled BACH2-high TRM Th1_1 and TRM Th17 cells, likely because the majority of CD4+ T cell clones were identified in these two clusters. Nevertheless, we determined that CD4+ T cell clones in PLWH were more activated (by antigen stimulation) when compared to CD4+ T cell clones in HIV– individuals, albeit PLWH participants were under suppressive ART.

### HIV-1-infected cells were enriched in gut TRM Th17 and TRM Th1

Beyond the current pursuit of the cell type and cellular markers of HIV-1-infected cells, we aim to interrogate how HIV-1-infected cells survive and persist in the gut. To examine the epigenetic, transcriptional, and protein expression differences between HIV-1-infected versus uninfected CD4+ T cells in PLWH under ART, we mapped ATAC-seq reads and RNA-seq reads to a compendium of clade B HIV-1 sequences to identify HIV-1 DNA+ cells and HIV-1 RNA+ cells (see Methods). In total, we identified 54 HIV-1 DNA+ RNA– cells (transcriptionally inactive HIV-1-infected cells) and 45 HIV-1 RNA+ cells (transcriptionally inactive HIV-1-infected cells) for a total of 99 HIV-1-infected cells from ten PLWH participants (**Figure 6A**). We defined HIV-1 DNA+ RNA– cells as HIV-1 DNA+ cells having undetectable HIV-1 expression. Of note, because of the sparsity of HIV-1 DNA captured in high-throughput single-cell short-read sequencing, this method cannot determine the intactness of HIV-1 DNA. HIV-1 DNA+ cells likely harbor defective proviruses^61^. We pooled HIV-1 DNA+ RNA– cells and HIV-1 RNA+ cells as HIV-1-infected cells to ensure statistical rigor for analysis. Per participant under suppressive ART, we found that 0.04% to 0.74% of gut CD4^+^ T cells were HIV-1-infected (**Figure S9A, Table S2, Table S5**). Most HIV-1 DNA reads were captured within the HIV-1 LTR promoter, reflecting increased chromatin accessibility at HIV-1 promoter than protein coding regions, whereas most HIV-1 RNA reads were captured across the genome, likely because of poly-T priming in multiple A-rich regions in HIV-1 genome (**Figure S9B, S9C**).

**Figure 6.**
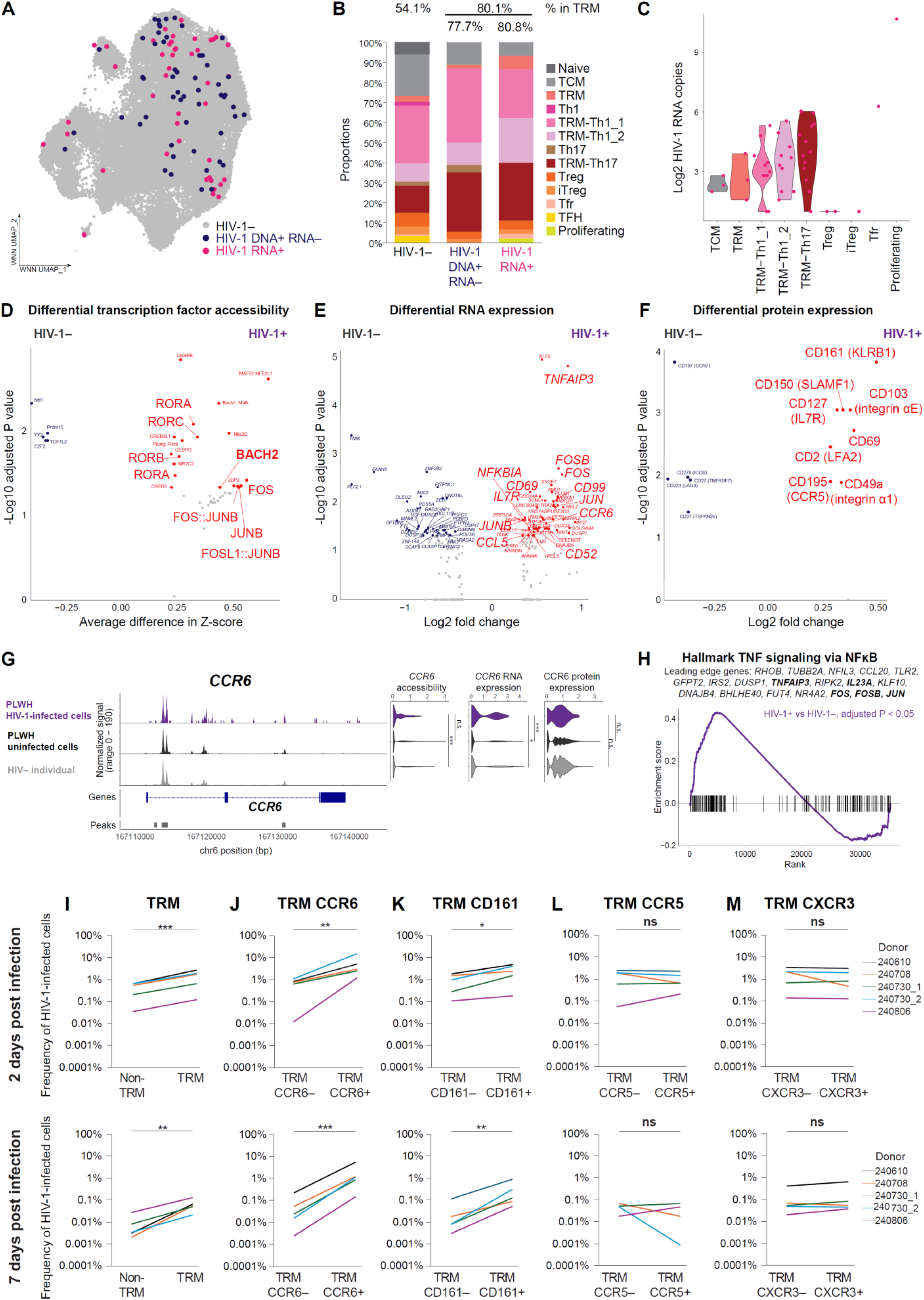
HIV-1 preferentially infects and persists in TRM CCR7+ KLRB1+ CD4+ T cells in the gut. (**A**) WNN UMAP plot depicting a total of 99 HIV-1-infected cells, including 54 HIV-1+ DNA+ RNA− cells (blue) and 45 HIV-1 RNA+ cells (magenta). (**B**) Cell subset proportions of HIV-1-infected cells. (**C**) Distribution of log-transformed HIV-1 RNA copies detected in HIV-1 RNA+ cells in each CD4+ T cell subset. (**D**) Differential transcription factors binding motifs accessibility (measured by chromVAR bias-corrected deviations) between HIV-1+ cells and HIV-1– CD4+ T cells in PLWH. FDR-adjusted *P* < 0.05, average Z-score difference > 0.25. (**E**) Differentially expressed genes (normalized and scaled) between HIV-1+ cells and HIV-1- CD4+ T cells in PLWH. FDR-adjusted *P* < 0.05, min.pct > 0.3, log2FC > 0.33. (**F**) Differentially expressed surface proteins (normalized and scaled) between HIV-1+ cells and HIV-1- CD4+ T cells in PLWH. FDR-adjusted *P* < 0.05, min.pct > 0.15, log2FC > 0.3. The mean expression of all protein features shown were tested to be greater than the mean expression of their specific isotype controls (*Z* > 2; two-sample Z test) in CD4+ T cells. For fair comparisons between groups, HIV-1− cells were downsized to match the same number of cells in HIV-1+ cells (n = 99) with 1,000 bootstrap replicates. The 1,000 *P* values were transformed to follow a standard Cauchy distribution and a combined *P* value was calculated as the weighted sum of transformed p values, followed by correction for multiple comparisons using the Benjamini-Hochberg (FDR) procedure. (**G**) Genome tracks showing *CCR6* gene locus chromatin accessibility, and violin plots showing average normalized and scaled chromatin accessibility at *CCR6* locus, *CCR6* gene expression, and CCR6 protein expression. (**H**) To identify enriched pathways in HIV-1+ cells, all 36,601 genes were ranked by log2-fold change in normalized gene expression between HIV-1+ cells and HIV-1- cells. Significantly enriched TNF signaling was identified in HIV-1+ cells by GSEA with leading-edge genes shown. **(I,J,K,L,M)** Flow cytometry of HIV-1-infected CD4+ TRM cells and HIV-1-infected CD4+ non-TRM cells at 2 days post infection (top) and 7 days post infection (bottom). The lamina propria layer of human colon was isolated, activated with CD3/CD28, and infected with HIV-dEnv-dNef-GKO pseudotyped with an R5 envelope (JR-FL). Each line represents a biological replicate from uninfected donors. TRM was defined as CD49+CD69+CD4+ T cells. ∗ p < 0.05, ∗∗ p < 0.01, ∗∗∗ p < 0.001, Wilcoxon rank-sum test. See also Figure S9 – S10.

We found that 80.80% of HIV-1-infected cells reside in TRMs, despite that TRMs only account for 54.11% of uninfected CD4+ T cells in PLWH (**Figure 6B**, **Table S2**). Enrichment of HIV-1-infected cells was most prominent in BACH2-high TRM Th17 cells (29.63% for HIV-1 DNA+ cells and 28.89% for HIV-1 RNA+ cells compared to 13.55% uninfected TRM Th17 cells in PLWH). Additionally, HIV-1 DNA+ cells were enriched in BACH2-high TRM Th1_1 (37.04% vs 28.80% uninfected cells in PLWH) but HIV-1 RNA+ cells were enriched in BACH2-low TRM Th1_2 (22.22% vs 9.06% uninfected cells in PLWH) (**Figure 6B, Table S2**). Furthermore, more HIV-1 RNA copies were identified from HIV-1 RNA+ cells in CD4+ TRMs (**Figure 6C**). Together, these results suggest that transcriptionally inactive (HIV-1 DNA+) HIV-1-infected cells were enriched in BACH2-high TRM Th1_1 and TRM Th17 that were long-lived memory TRMs with restrained effector T cell functions, whereas transcriptionally active (HIV-1 RNA+) cells were also enriched in TRM Th17 in addition to BACH2-low, short-lived, TRM Th1_2 cells that maintained effector Th1 functions. Overall, HIV-1-infected cells were considerably enriched in tissue resident memory CD4+ TRMs (80.81%).

### BACH2-driven long-lived tissue resident memory program promotes HIV-1 persistence

Comparing the chromatin accessibility profiles of HIV-1-infected cells vs uninfected cells in PLWH, we found that HIV-1-infected cells had increased transcription factor motif accessibility for AP-1 family, BACH2, and RAR orphan receptors (RORα, RORβ, and RORγt) (**Figure 6D**). Comparing the cellular transcriptional profiles of HIV-1-infected versus uninfected cells in PLWH, we found that HIV-1-infected cells had increased expression of chemokine *CCL5*, NF-κB inhibitor *NFKBIA*, chemokine receptors *IL7R* and *CCR6*, activation genes (*CD52*, *CD69*, *CD99*, *JUN*, *FOS*), and TNF response gene *TNFAIP3* (**Figure 6E**). Comparing the cell surface protein profiles of HIV-1-infected versus uninfected cells in PLWH, we found that HIV-1-infected cells had higher levels of markers of tissue resident adhesion molecules [CD103 (ITGAE), CD69, and CD49a (ITGA1)], Th17 marker CD161, chemokine receptors CCR5 and cytokine receptor IL7R, and activation marker CD150 (SLAMF1), (**Figure 6F**). Overall, comparing HIV-1-infected cells to uninfected cells in PLWH, we found that the cellular profiles of HIV-1-infected cells closely resembled that of BACH2-driven long-lived TRMs, consistent with the finding that 60.61% of all HIV-1-infected cells resided in BACH2-high CD4+ TRM Th1_1 and TRM Th17 (**Table S2**). To mitigate noise in cellular profile of HIV-1-infected cell introduced by cell type bias in TRMs, we repeated differential expression analyses for HIV-1-infected TRMs vs uninfected TRMs in PLWH. We found that HIV-1-infected TRMs had increased transcription factor accessibility for interferon regulatory factor IRF3 and RORA (Th17) (**Figure S9D**), higher RNA expression of Th17 marker *CCR6*, activation marker *ITM2A*, and TNF-induced *TNFAIP3* gene expression (**Figure S9E**), and increased protein expression of tissue resident markers CD103 (ITGAE) and CD69, activation CD150 (SLAMF1), CD161 (KLRB1), and CD28 (**Figure S9F**). Indeed, Th17 gut-homing receptor *CCR6*^62^ (**Figure 6G**), tissue resident *ITGAE*^50,51^ (**Figure S9G**), and long-lived memory *IL7R*^63^ (**Figure S9H**) had increased chromatin accessibility at gene locus, and upregulated RNA and protein expression, in HIV-1-infected cells.

Lastly, to determine enriched gene pathways in HIV-1-infected cells vs uninfected cells in PLWH, Gene Set Enrichment Analysis (GSEA) revealed significant enrichment in TNF signaling response (leading edge genes *TNFAIP3*, *TNIP2*, *CCL20*, *IL15RA*, *IL23A*, *FOS*, *JUN*) in HIV-1-infected cells (**Figure 6H**), and KEGG pathway analysis revealed significant enrichment of TNF signaling pathway, IL17 signaling pathway, Th1 and Th17 differentiation, and TCR signaling pathway in HIV-1-infected cells (**Figure S9I**). Overall, we found that HIV-1-infected cells demonstrated strong responses in tissue residency, Th17 differentiation, and TNF responses.

### HIV-1 preferentially infects and persists in CCR6+ KLRB1+ TRM

To validate whether HIV-1 preferentially infects and persists CD4+ TRM, we infected primary cells isolated from the lamina propria of colon excisions from four uninfected donors (**Table S1**) with a single-round HIV-dEnv-dNef-GKO reporter^64^ pseudotyped with a R5 envelope (JR-FL) after CD3/CD28 activation. This HIV-1 reporter contains intact Vpr, Vpu, and Vif and can cause viral cytopathic effects in infected cells^65^, but does not have functional Nef and thus does not effectively downregulate CD4 expression. We examine whether HIV-1 preferentially infects CD4+ TRM by measuring the proportion of HIV-1-infected cells 2 days after infection, and whether HIV-1 preferentially persists in CD4+ TRM by measuring the proportion of HIV-1-infected cells 7 days after infection, when viral cytopathic effect should have killed most infected cells (**Figure S10A**). CD4+ cells expressing both CD49a (integrin α1) and CD69 were defined as TRMs. We found that the proportion of HIV-1-infected TRMs was significantly higher than those in non-TRMs at both 2 and 7 days after infection, indicating preferential HIV-1 infection and persistence in TRMs (**Figure 6I**). We next tested whether HIV-1 preferentially infects or persists in cells expressing Th17 markers (CCR6 and CD161) and Th1 markers (CCR5 and CXCR3). We found that the proportion of HIV-1-infected cells was higher in CCR6+ TRMs (**Figure 6J**) and CD161+ TRMs (**Figure 6K**) both 2 days and 7 days after infection, indicating preferential infection and persistence of HIV-1 infection in CCR6+ TRMs and CD161+ TRMs. Surprisingly, although HIV-1 infection is thought to be enriched in CCR5+CD4+ T cells^66–68^, we found that the proportion of HIV-1-infected cells in CCR5+ TRMs and CXCR3+ TRMs was not higher than CCR5– TRMs (**Figure 6L**) and CXCR3– TRMs (**Figure 6M**). Of note, the proportion of HIV-1-infected cells in CCR6+ CD4+ T cells and CD161+ CD4+ T cells (including both TRMs and non-TRMs) was still higher than those in CCR6– CD4+ T cells and CD161– CD4+ T cells (**Figure S10B, S10C**), indicating CCR6 and CD161 were markers for HIV-1-infected CD4+ T cells in the gut. Proportions of HIV-1-infected cells were neither higher in CCR5+ CD4+ T cells (**Figure S10D**) nor in CXCR3+ CD4+ T cells (**Figure S10E**). Together, the highest-level enrichment of HIV-1-infected cells was found in CCR6+ Th17 (**Figure 6J**). Our *in vitro* validation of HIV-1 infection of primary gut CD4+ T cells showed that HIV-1 preferentially infects and persists in TRM Th17, which is consistent with our single-cell multiomic analysis (**Figure 6D – 6F**).

## DISCUSSION

The combination of single-cell multiomic profiling and gene regulatory network analysis identified BACH2 as the key transcription factor in the gut, which drives both CD4+ (**Figure 1F**) and CD8+ (**Figure 2C**) TRMs into long-lived memory cells. Although it takes weeks to develop adaptive immune response upon the first antigen encounter, these long-lived memory T cells can respond to specific antigen challenge rapidly to provide immediate local responses without systemic inflammation. BACH2-driven tissue resident program retains long-lived memory T cells locally at the mucosal site, restricts excessive effector responses, and allows persistence of pathogen-specific memory within the gut. HIV-1 infects and depletes gut CD4+ T cells, disrupts mucosal barrier, and increases microbial translocation^69^. Presumably, the most important effector T cells in PLWH gut are TRMs, including TRM Th1 that respond to HIV-1 infection, TRM Th17 that repair and seal intestinal epithelial tight junctions (by IL-17 induced upregulation of occludin^70^) and reduce microbial translocation, and cytotoxic CD8+ T cells that eliminate HIV-1-infected cells.

We found that BACH2-driven tissue residency program promoted HIV-1 persistence. While IRF drove effector T cell responses (**Figure 3**) and CD8+ T cell proliferation (**Figure 5**), BACH2-driven tissue resident program attenuated effector responses and increased cytokine receptor expression to maintain long-term survival (**Figure 3**). BACH2-low CD8+ T cells, while exerted effector function by IFN-γ and TNF secretion, promoted the survival of BACH2-high TRM Th1 and TRM Th17 (as shown by *BCL2* and *IL7R* expression, **Figure 4E, 4F**) by IFN-γ and TNF ligand-receptor interactions (**Figure 4C, 4D**). The long-lived nature of CD4+ TRMs allowed HIV-1-infected cells to upregulate survival cytokine receptors (**Figure 6**), escape activation-induced cell death, and persist in the gut.

Our study identified several CD8+ T cell populations that may exert cytotoxic potential, including CD8+ TEM (**Figure 2**), BACH2-low CD8 TRM_1 (**Figure 3**), and CD8+ T cell clones (**Figure 5**), as evidenced by high RNA expressions of GZMA molecules^71–74^. Indeed, intestinal intraepithelial T lymphocytes (IELs) mediate GZMA-driven apoptosis that effectively reduced Salmonella growth and translocation across the intestinal barrier^60^. Nevertheless, the cytolytic potential of CD8+ T cells in tissues is debated^60,75,76^ and granzyme molecules may also induce caspase-independent inflammation to exert tissue damage.

Under ART, HIV-1-specific CD8+ T cells exhibited CD8+ TRM phenotype, low proliferation, and reduced effector function (**Figure S6**). HIV-1-induced chronic inflammation in the gut led to clonal proliferation of CD8+ T cells that was far greater than CD4+ T cells (**Figure 5A**). These CD8+ T cell clones were signatured with increased IRF transcription factor activity and increased effector functions (**Figure 5I**). However, the vast majority of CD8+ T cell clones were bystander and not HIV-1-specific. Consistent with a previous study^77^, we found that most HIV-1-specific CD8+ T cells were TRMs. Because BACH2-driven TRM program maintained long-lived memory cells and restrained effector T cell function, TRM-enriched HIV-1-specific CD8+ T cells had attenuated effector function in comparison to CMV-specific or bystander CD8+ T cells that were not TRM-specific (**Figure S6**). Nonetheless, we found that HIV-1-specific CD8+ T cells were not exhausted, likely because ART rejuvenated these cells^31^. Under ART, HIV-1-specific CD8+ T cells declined but persisted in TRM and had reduced effector functions that hampered HIV-1 immune clearance. Our results highlight the need of reprogramming dysfunctional HIV-1-specific CD8+ T cells back to functional effectors in the gut.

Our study revealed the dramatic difference in mechanisms of HIV-1 persistence in the gut versus the existing knowledge based on blood immune cell profiling. HIV-1 is known to preferentially infect and persist in Th1 and Th17 cells in blood^15,17,78,79^, CCR6+ CD4+ T cells in the gut^22,80,81^, and in CD4+ TRMs in cervical tissue^82^. However, how HIV-1 persists in the gut remains unknown. We postulated that HIV-1 persists in the gut by residing in BACH2-high CD4+ TRMs, as these cells persist long-term, have restrained effector functions and attenuated activation-induced cell death. Indeed, HIV-1-infected cells were enriched in CD4+ TRMs (80.81%), given that TRMs in uninfected cells accounted only around half (54.11%) of gut CD4+ T cells. Contrary to common belief that HIV-1 persists in Tbet-driven Th1 CD4+ T cells expressing high levels of CCR5^66–68^ or cytotoxic CD4+ T cells^15^, we found that HIV-1-infected cells in the gut exhibit TRM Th17 phenotype as CCR6+ TRMs and CD161+ TRMs, as shown at the epigenetic level (increased BACH2, AP-1, and RORC transcription factor accessibility, **Figure 6D**), transcriptome level (*CCR6*, *IL7R*, and *TNFAIP3* gene expression, **Figure 6E**), and protein expression level (CD49a, CD69, and CD161 protein expression, **Figure 6F**). We speculate that with constant antigen stimulation of Th17, likely from microbial translocation, the maintenance of TRM Th17 promotes the survival of HIV-1-infected TRM Th17 (**Figure 6G, 6I – 6K**).

Our study indicates that HIV-1 infection in high BACH2 transcription factor activity, long-lived tissue resident memory, CD4+ T cells allowed infected cells to persist. Whether BACH2 as a transcription repressor suppresses HIV-1 promoter activity and thus reduces reactivation and immune recognition remains to be examined. Our study identifies BACH2 as a potential therapeutic target for eradicating HIV-1 gut reservoir in gut TRMs to accelerate HIV-1 eradication.

### Limitations of the Study

The major limitations of the study were the low number of HIV-1-infected CD4+ T cells and HIV-1-specific CD8+ T cells identified from PLWH under suppressive ART which limited the conclusions that could be drawn about these cells, and the inability to infer proviral genome intactness (DOGMA-seq recovers fragmented DNA and RNA reads). Additionally, the TCR capturing efficiency (percentage of cells with TCR determined) using the 3’-based TREK-seq method is less than the 5’-based method in ECCITE-Seq that we previously reported^15^, however no other current methods enable sequencing of ATAC, RNA, barcoded surface protein, and TCR in the same single cells. Lastly, nine out of ten PLWH studied were male, so the generalizability of our conclusions to people of different sex remains to be determined, although part of our results have been confirmed by *in vitro* validation in female participants.

## AUTHOR CONTRIBUTIONS

Y.W. and Y.-C.H. designed and performed DOGMA-seq, TREK-seq, and tissue isolation experiments. Y.W. performed bioinformatic analyses. H.K.M performed flow cytometry. M.E.W. designed tissue isolation protocol. L.K. reviewed the study design and processed clinical samples. P.T. and R.M. recruited study participants and processed clinical samples. E.P. and L.J.M. designed and conducted the clinical study. Y.W. and Y.-C.H. wrote the manuscript with input from all authors.

## SUPPLEMENTARY FIGURE LEGENDS

**Figure S1. Identification of surface protein epitopes not degraded by collagenase treatment, refers to Figures 1, 2, 5, and 6.** (**A**) Heatmap indicating average CLR-normalized and scaled expressions of all 155 surface proteins in TotalSeq-A cocktail plus TotalSeq-A CD197 (CCR7), in 3 mock (untreated) and 3 collagenase II treated PBMC aliquots from an uninfected donor. (**B**) Distributions of CLR-normalized and scaled protein expression of CD183 [CXCR3], CD185 [CXCR5], CD45RA, CD56, CD119 [IFNGR1], and CD4 in mock and collagenase II treated PBMC. From each pooled sample, cells with high protein expressions for each protein (> 95^th^ percentile) were considered for comparisons to mitigate noise from ambient background expression. (**C**) Heatmap indicating average CLR-normalized and scaled expressions of all 155 surface proteins in TotalSeq-A cocktail plus TotalSeq-A CD197 (CCR7), in 3 mock and 3 collagenase IV treated PBMC aliquots from the same uninfected donor. ∗ *P* < 0.05, Wilcoxon rank-sum test.

**Figure S2. Identities of gut CD4+ T cell subsets as defined by epigenetic, transcriptional, and surface protein states, refers to Figure 1**. (**A**) WNN UMAP plot showed integration of CD4+ T cells (n = 49,205) across samples from PLWH and HIV– individuals following batch effect removal by Harmony (RNA) and reciprocal LSI (ATAC). (**B,C,D**) Key epigenetic, transcriptional, and protein markers used to manually annotate all 13 computationally identified CD4+ T cell clusters. (**B**) Average normalized and scaled chromatin accessibility (ATAC) at gene locus of marker genes. (**C**) Average normalized and scaled RNA expression. (**D**) Average CLR-normalized and scaled surface protein expression. To mitigate noise from ambient background expression, for each protein, only cells having the highest expressions (expression > 95^th^ percentile of the expression level of specific isotype controls and >95^th^ percentile expression in all cells) were used to determine percentage of cells stained per cell cluster. (**E**) To identify enriched pathways in CD4+ TRM T cells, all 36,601 genes were ranked by log2-fold change in normalized gene expression between CD4+ TRM cells and non-TRM CD4+ cells. Significantly enriched IFNγ response was identified in PLWH TRM and HIV– individuals TRM cells using GSEA with top leading-edge genes shown.

**Figure S3. Characteristics of BACH2-driven gut CD4+ tissue resident memory program, refers to Figure 1**. (**A**) Heatmap of DORC regulation scores for all significant TF-DORC associations identified by GRN, for DORC genes that were upregulated in CD4+ T cells in PLWH (FDR-adjusted *P* < 0.05, minimum percentage of cells > 0.1, log_2_ fold change > 0.1). (**B**) BACH2 transcription factor accessibility in CD4+ T cells, measured by chromVAR bias-corrected deviations. (**C**) RNA module score of DORC genes (upregulated in PLWH and predicted to be activated by BACH2 in CD4+ T cells). (**D**) Cnetplot, depicts top enriched Gene Ontology Biological Processes (GO:BP) pathways (blue) and genes related to pathways (black) determined using DORC genes upregulated in PLWH and predicted to be activated by BACH2 in CD4+ T cells. (**E,F,G**) Genome tracks showing gene locus chromatin accessibility, and dot plots showing average normalized and scaled gene accessibility and RNA expression, for *BACH2*, *ITGA1*, *ITGAE*, *CD69* (**E**), *TBX21*, *CXCR3*, *CCR5*, *CCL5* (**F**), and *RORC*, *CCR6*, *CCL20*, *KLRB1* (**G**). Highlighted red on the genome track corresponds to significantly differentially accessible peaks called by MACS2 (FDR-adjusted *P* < 0.05, minimum percentage > 0.01, log2 fold change > 0.1). ∗ *P* < 0.05, ∗∗ *P* < 0.01, ∗∗∗ *P* < 0.001, Wilcoxon rank-sum test.

**Figure S4. Identities of gut CD8+ T cell subsets as defined by epigenetic, transcriptional, and surface protein states, refers to Figure 2**. (**A**) WNN UMAP plot showed integration of CD8+ T cells (n = 51,539) across samples from PLWH and HIV– individuals following batch effect removal by Harmony (RNA) and reciprocal LSI (ATAC). (**B,C,D**) Key epigenetic, transcriptional, and protein markers used to manually annotate all 11 computationally identified cell clusters. (**B**) Dot plot showed average normalized and scaled chromatin accessibility (ATAC) at gene locus of marker genes. (**C**) Average normalized and scaled RNA expression. (**D**) Average CLR-normalized and scaled surface protein expression. To mitigate noise from ambient background expression, for each protein, only cells having the highest expressions (expression > 95^th^ percentile of the expression level of specific isotype controls and >95^th^ percentile expression in all cells) were used to determine percentage of cells stained per cell cluster. (**E**) Heatmap of gene regulation scores for all significant TF to gene associations identified by GRN, for DORC genes that were upregulated in CD8+ T cells in PLWH (FDR-adjusted *P* < 0.05, minimum percentage > 0.1, log_2_ fold change > 0.1).

**Figure S5. BACH2 restrains effector T cell programs in TRM cells that had enriched TNF and IFN-γ signaling responses, refers to Figures 3 and 4**. (**A, B**) Dynamic IRF transcription factor accessibility changes with increase in BACH2 transcription factor accessibility. Accessibility of IRF family transcription factor binding motifs decreased with increase in BACH2 motif accessibility in CD4+ TRM Th1 cells (TRM Th1_1 and TRM Th1_2 combined) (**A**) and in CD8+ TRM cells (**B**). (**C, D**) Scriabin identified 2 statistically significant ligand – receptor interaction programs, IP-2 (**C**) and IP-12 (**D**), between sender activated CD8+ T cells and TRM Th1_2 cells (blue) and receiver TRM Th1_1 and TRM Th17 cells (red). (**E, F**) Intramodular connectivity of all ligand – receptor pairs identified in IP-2 (**E**) and IP-12 (**F**), in T cells from PLWH and HIV– individuals. (**G, H**) Dot plot depicting NicheNet-predicted ligand activity in receiver TRM-Th1_1 (**G**) and TRM Th17 (**H**) in PLWH and HIV– individuals. Dot size depict percentage of receiver cells having ligand activity weight of > 0.1. Color scales represent ligand expression from CD8 activated sender cells (measured by RNA expression fold change in CD8 activated relative to other cell clusters).

**Figure S6. HIV-1 antigen-specific CD8+ T cells exhibit tissue resident phenotype and restrained cytotoxic effector molecule expression, refers to Figure 5**. (**A**) WNN UMAP plot of 10,868 CD8+ T cells whose TCRβ CDR3 junction sequences (CDR3β) were captured, including 56 predicted HIV-1-specific (red) and 429 CMV-specific (blue) CD8+ T cells. (**B**) Cell subset proportions of HIV-1-specific, CMV-specific, and other CD8+ T cells with mapped TCRβ. (**C**) Differentially expressed genes (normalized and scaled) between HIV-1-specifc CD8+ T cell and CMV-specific CD8+ T cells. FDR-adjusted *P* < 0.05, min.pct > 0.25, log2FC > 0.5. (**D**) Differentially expressed genes (normalized and scaled) between HIV-1-specifc CD8+ T cell and other CD8+ T cells. FDR-adjusted *P* < 0.05, min.pct > 0.25, log2FC > 0.5. (**E**) Differentially expressed surface proteins (normalized and scaled) between HIV-1-specifc CD8+ T cell and CMV-specific CD8+ T cells. FDR-adjusted *P* < 0.05, min.pct > 0.1, log2FC > 0.25. (**F**) Differentially expressed surface proteins (normalized and scaled) between HIV-1-specifc CD8+ T cell and other CD8+ T cells. FDR-adjusted *P* < 0.05, min.pct > 0.1, log2FC > 0.25. The mean expression of all protein features shown were tested to be greater than the mean expression of their specific isotype controls (*Z* > 2; two-sample Z test) in CD8+ T cells. (**G,H**) Differences in averaged and scaled RNA (**G**) and protein (**H**) expression of select features between HIV-1-specific, CMV-specific, and other CD8+ T cells. (**I**) Alluvial plot indicating the proportions of CD8+ T cell clones by shared CDR3β junction sequences in each participant (indicated by bar size), the size of each clone per participant (indicated by bar size), and antigen-specificity for each clone (HIV-1 antigen-specific in red, CMV antigen-specific in blue).

**Figure S7. Epigenetic, transcriptional, and surface protein states of CD8+ T cell expanded clones, refers to Figure 5**. (**A**) Differential transcription factors binding motif accessibility (measured by chromVAR bias-corrected deviations) between CD8+ TRM clones and cells not clonal. FDR-adjusted *P* < 0.05, average Z-score difference > 0.1. (**B**) Differentially expressed genes (normalized and scaled) between CD8+ TRM clones and CD8+ TRM cells not clonal. FDR-adjusted *P* < 0.05, min.pct > 0.25, log2FC > 0.25. (**C**) Differentially expressed surface proteins (normalized and scaled) between CD8+ TRM clones and CD8+ TRM cells not clonal. FDR-adjusted *P* < 0.05, min.pct > 0.1, log2FC > 0.1. The mean expression of all protein features shown were tested to be greater than the mean expression of their specific isotype controls (*Z* > 2; two-sample Z test) in tissue-resident CD8+ TRM cells. For all comparisons, only CD8+ TRM cells having identified CDR3β were used. (**D**) Differential transcription factors binding motif accessibility (measured by chromVAR bias-corrected deviations) in CD8+ T cell clones between PLWH and HIV– individuals. FDR-adjusted *P* < 0.05, average Z-score difference > 0.1. (**E**) Differentially expressed genes (normalized and scaled) in CD8+ T cell clones between PLWH and HIV– individuals. FDR-adjusted *P* < 0.05, min.pct > 0.25, log2FC > 0.25. (**F**) Differentially expressed surface proteins (normalized and scaled) in CD8+ T cell clones between PLWH and HIV– individuals. FDR-adjusted *P* < 0.05, min.pct > 0.1, log2FC > 0.25. The mean expression of all protein features shown were tested to be greater than the mean expression of their specific isotype controls (*Z* > 2; two-sample Z test) in CD8+ T cell clones. For fair comparisons between PLWH and HIV– individuals CD8+ T cell clones, PLWH CD8+ T cell clones were downsized to match the same number of cells in HIV-1+ cells (n = 308) with 1,000 bootstrap replicates. The 1,000 *P* values were transformed to follow a standard Cauchy distribution and a combined *P* value is calculated as the weighted sum of transformed *P* values, followed by correction for multiple comparisons using the Benjamini-Hochberg (FDR) procedure. (**G,H**) Genome tracks showing gene locus chromatin accessibility, and dot plots showing average normalized and scaled chromatin accessibility at gene locus and RNA expression, for *CCL5* and *GZMA* genes that were upregulated in RNA expression in PLWH CD8+ T cell clones (**G**) and for *BACH2* and *CCR7* genes that were downregulated in RNA expression in PLWH CD8+ T cell clones (**H**) in comparison to HIV– individuals CD8+ T cell clones. Highlighted red on the genome track corresponds to significantly differentially accessible peaks called by MACS2 (FDR-adjusted *P* < 0.05, min.pct > 0.01, log2FC > 0.1). (**I,J**) Top enriched Gene Ontology Biological Processes (GO:BP) pathways, using genes upregulated in PLWH CD8+ T cell clones vs PLWH CD8+ T cell not clonal (**I**), and using genes upregulated in PLWH CD8+ T cell clones vs HIV– individuals CD8+ T cell clones (**J**). ∗ *P* < 0.05, ∗∗ *P* < 0.01, ∗∗∗ *P* < 0.001, Wilcoxon rank-sum test.

**Figure S8. Epigenetic, transcriptional, and surface protein states of CD4+ T cell expanded clones, refers to Figure 5**. (**A**) Differential transcription factors binding motif accessibility (measured by chromVAR bias-corrected deviations) between CD4+ T cell clones and CD4+ T cells not clonal in PLWH. FDR-adjusted *P* < 0.05, average Z-score difference > 0.1. (**B**) Differentially expressed genes (normalized and scaled) between CD4+ T cell clones and CD4+ T cells not clonal in PLWH. FDR-adjusted *P* < 0.05, min.pct > 0.25, log2FC > 0.25. (**C**) Differentially expressed surface proteins (normalized and scaled) between CD4+ T cell clones and CD4+ T cells not clonal in PLWH. FDR-adjusted *P* < 0.05, min.pct > 0.25, log2FC > 0.25. (**D**) Differential transcription factors binding motif accessibility (measured by chromVAR bias-corrected deviations) between CD4+ TRM clones and CD4+ TRM cells not clonal in PLWH. FDR-adjusted *P* < 0.05, average Z-score difference > 0.1. (**E**) Volcano plots indicating differentially expressed genes (normalized and scaled) between CD4+ TRM clones and CD4+ TRM cells not clonal in PLWH. FDR-adjusted *P* < 0.05, min.pct > 0.25, log2FC > 0.25. (**F**) Volcano plots indicating differentially expressed surface proteins (normalized and scaled) between CD4+ TRM clones and CD4+ TRM cells not clonal in PLWH. FDR-adjusted *P* < 0.05, min.pct > 0.25, log2FC > 0.25. For all comparisons, only CD4+ cells having identified CDR3β were used. (**G**) Differential transcription factors binding motif accessibility (measured by chromVAR bias-corrected deviations) between CD4+ T cell clones in PLWH and CD4+ T cell clones in HIV– individuals. FDR-adjusted *P* < 0.05, average Z-score difference > 0.1. (**H**) Differentially expressed genes (normalized and scaled) between CD4+ T cell clones in PLWH and CD4+ T cell clones in HIV– individuals. FDR-adjusted *P* < 0.05, min.pct > 0.25, log2FC > 0.25. (**I**) Volcano plots indicating differentially expressed surface proteins (normalized and scaled) between CD4+ T cell clones in PLWH and CD4+ T cell clones in HIV– individuals. FDR-adjusted *P* < 0.05, min.pct > 0.25, log2FC > 0.25. The mean expression of all protein features shown were tested to be greater than the mean expression of their specific isotype controls (*Z* > 2; two-sample Z test) in CDR3β-captured CD4+ T cells. For fair comparisons between groups, cells were downsized to match the same number of cells in both conditions with 1,000 bootstrap replicates. The 1,000 *P* values were transformed to follow a standard Cauchy distribution, and a combined p value is calculated as the weighted sum of transformed p values, followed by correction for multiple comparisons using the Benjamini-Hochberg (FDR) procedure.

**Figure S9. HIV-1-infected cells upregulate TRM Th17 genes, refers to Figure 6**. (**A**) Per participant distribution of HIV-1+ DNA+ RNA− cells (blue) and HIV-1 RNA+ cells (magenta) visualized in WNN UMAP projection. (**B,C**) HIV-1 proviral and viral genome landscape captured by DOGMA-seq. Integrative genomics viewer (IGV) plots of HIV-1 DNA^+^ reads (top) and of HIV-1 RNA^+^ reads (bottom) that mapped to HXB2 reference genome and Clade B compendium reference sequences, in all PLWH CD4+ T cells (**B**) and in per PLWH participant CD4+ T cells (**C**). (**D**) Differential transcription factors binding motif accessibility (measured by chromVAR bias-corrected deviations) between HIV-1+ TRM cells and HIV-1-CD4+ TRM T cells in PLWH. FDR-adjusted *P* < 0.05, average Z-score difference > 0.2. (**E**) Volcano plots showing differentially expressed genes (normalized and scaled) between HIV-1+ TRM cells and HIV-1-CD4+ TRM T cells in PLWH. FDR-adjusted *P* < 0.05, min.pct > 0.33, log2FC > 0.44. (**F**) Differentially expressed surface proteins (normalized and scaled) between HIV-1+ TRM cells and HIV-1-CD4+ TRM T cells in PLWH. FDR-adjusted *P* < 0.05, min.pct > 0.15, log2FC > 0.3. The mean expression of all protein features shown were tested to be greater than the mean expression of their specific isotype controls (*Z* > 2; two-sample Z test) in CD4+ TRM T cells. For fair comparisons between groups, HIV-1− TRM T cells were downsized to match the same number of cells in HIV-1+ cells (n = 99) with 1,000 bootstrap replicates. The 1,000 *P* values were transformed to follow a standard Cauchy distribution and a combined p value is calculated as the weighted sum of transformed p values, followed by correction for multiple comparisons using the Benjamini-Hochberg (FDR) procedure. (**G,H**) Genome tracks showing gene locus chromatin accessibility, and violin plots showing average normalized and scaled chromatin accessibility, gene expression, and protein expression, for *ITGAE* (**G**) and for *IL7R* (**H**). Highlighted red on the genome track corresponds to significantly differentially accessible peaks called by MACS2 (FDR-adjusted *P* < 0.05, min.pct > 0.01, log2FC > 0.1). **i.** KEGG pathways enriched in HIV-1+ cells determined using genes upregulated in HIV-1+ cells compared to HIV-1-CD4+ T cells. ∗ *P* < 0.05, ∗∗ *P* < 0.01, ∗∗∗ *P* < 0.001, Wilcoxon rank-sum test.

**Figure S10. Gating strategies of in vitro validation of HIV-1-infection in primary gut cells, refers to Figure 6**. (**A**) Representative flow cytometry plot showing gating strategies of HIV-1-infected tissue resident memory (CD69+ CD49a+, double positive) CD4+ T cells, and of CCR6+, CD161+, CCR5+, and CXCR3+ HIV-1-infected cells. Lamina propria layer of gut cells were isolated, activated with CD3/CD28, and infected with HIV-1-dEnv-dNef-GKO pseudotyped with an R5-tropic (JR-FL) envelope. (**B,C,D,E**) Flow cytometry of HIV-1-infected gut CD4+ T cells at 2 days post infection (top) and 7 days post infection (bottom). Each line represents a biological replicate from uninfected donors. ∗ *P* < 0.05, ∗∗ *P* < 0.01, ∗∗∗ *P* < 0.001, Wilcoxon rank-sum test.

## Supporting information

2024 Yulong Wei HIV gut DOGMAseq supplement

## ACKNOWLEDGEMENTS

We thank all study participants. We thank Kenneth Lynn and Beth Peterson for supporting study participant recruitment. We thank Guilin Wang and Yale Center for Genome Analysis. We thank Yalai Bai and Dongqing Liu from Yale Tissue Services, Department of Pathology and Oluwabunmi Olaloye for tissue processing guidance. We thank Joseph Craft for immunology insight and Cara Wilson, Mario Santiago, Stephanie Dillon, and Marcus Buggert for gut isolation advice. This work is supported by NIH R01 AI174863, R01 AI183430, R01 AI141009, R01 AI141009, P01 AI169768, NIH BEAT-HIV Martin Delaney Collaboratory UM1 AI164570, UM1 AI164565, CHEETAH U54 AI170856, R01 AI176601, R33 DA047037, UM1 M-SCORCH DA051410, U01 Y-SCORCH DA053628, R01 DA051906 (Y.-C.H.), Natural Sciences and Engineering Research Council of Canada (NSERC) Postdoctoral Fellowship (PDF557234) and AIDS and Cancer Specimen Resource Pilot Grant (Y.W.), and Yale Gruber Fellowship (H.K.M.).

