## Supplementary material for "BACH2-driven tissue resident memory programs promote HIV-1 persistence": 2024 Yulong Wei HIV gut DOGMAseq supplement

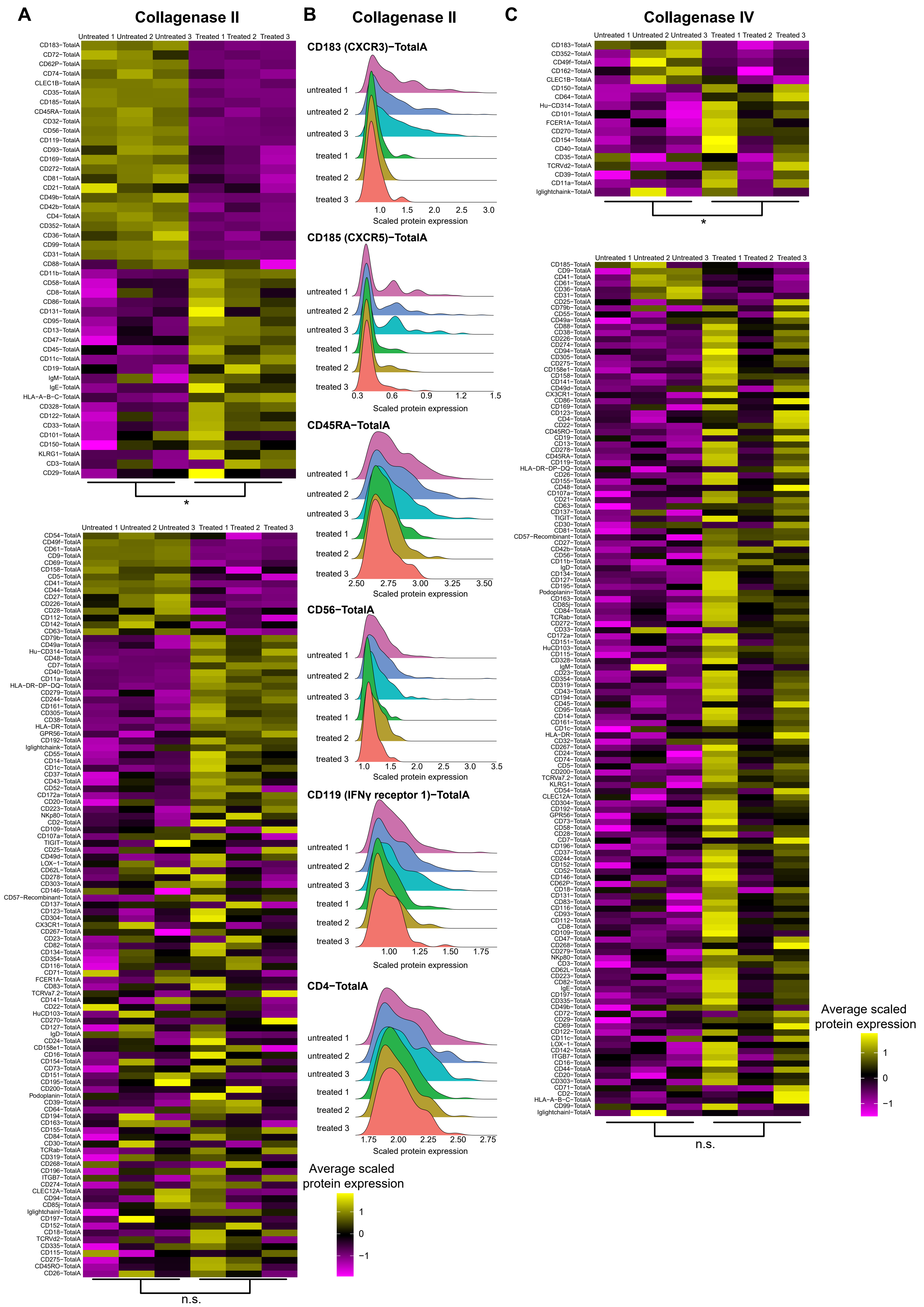
A
Collagenase II
Untreated 1 Untreated 2 Untreated 3 Treated 1 Treated 2 Treated 3
CD183−TotalA
CD72−TotalA
CD62P−TotalA
CD74−TotalA
CLEC1B−TotalA
CD35−TotalA
CD185−TotalA
CD45RA−TotalA
CD32−TotalA
CD56−TotalA
CD119−TotalA
CD93−TotalA
CD169−TotalA
CD272−TotalA
CD81−TotalA
CD21−TotalA
CD49b−TotalA
CD42b−TotalA
CD4−TotalA
CD352−TotalA
CD36−TotalA
CD99−TotalA
CD31−TotalA
CD88−TotalA
CD11b−TotalA
CD58−TotalA
CD8−TotalA
CD86−TotalA
CD131−TotalA
CD95−TotalA
CD13−TotalA
CD47−TotalA
CD45−TotalA
CD11c−TotalA
CD19−TotalA
IgM−TotalA
IgE−TotalA
HLA−A−B−C−TotalA
CD328−TotalA
CD122−TotalA
CD33−TotalA
CD101−TotalA
CD150−TotalA
KLRG1−TotalA
CD3−TotalA
CD29−TotalA
*
Untreated 1 Untreated 2 Untreated 3 Treated 1 Treated 2 Treated 3
n.s.
B
Collagenase II
CD183 (CXCR3)−TotalA
untreated 1
untreated 2
untreated 3
treated 1
treated 2
treated 3
1.0 1.5 2.0 2.5 3.0
Scaled protein expression
CD185 (CXCR5)−TotalA
0.3 0.6 0.9 1.2 1.5
Scaled protein expression
CD45RA−TotalA
2.50 2.75 3.00 3.25 3.50
Scaled protein expression
CD56−TotalA
1.0 1.5 2.0 2.5 3.0 3.5
Scaled protein expression
CD119 (IFNγ receptor 1)−TotalA
1.00 1.25 1.50 1.75
Scaled protein expression
CD4−TotalA
1.75 2.00 2.25 2.50 2.75
Scaled protein expression
Average scaled protein expression
1
0
−1
C
Collagenase IV
Untreated 1 Untreated 2 Untreated 3 Treated 1 Treated 2 Treated 3
CD183−TotalA
CD352−TotalA
CD49f−TotalA
CD162−TotalA
CLEC1B−TotalA
CD150−TotalA
CD64−TotalA
Hu−CD314−TotalA
CD101−TotalA
FCER1A−TotalA
CD270−TotalA
CD154−TotalA
CD40−TotalA
CD35−TotalA
TCRVd2−TotalA
CD39−TotalA
CD11a−TotalA
Iglightchaink−TotalA
*
Untreated 1 Untreated 2 Untreated 3 Treated 1 Treated 2 Treated 3
n.s.
Average scaled protein expression
1
0
−1

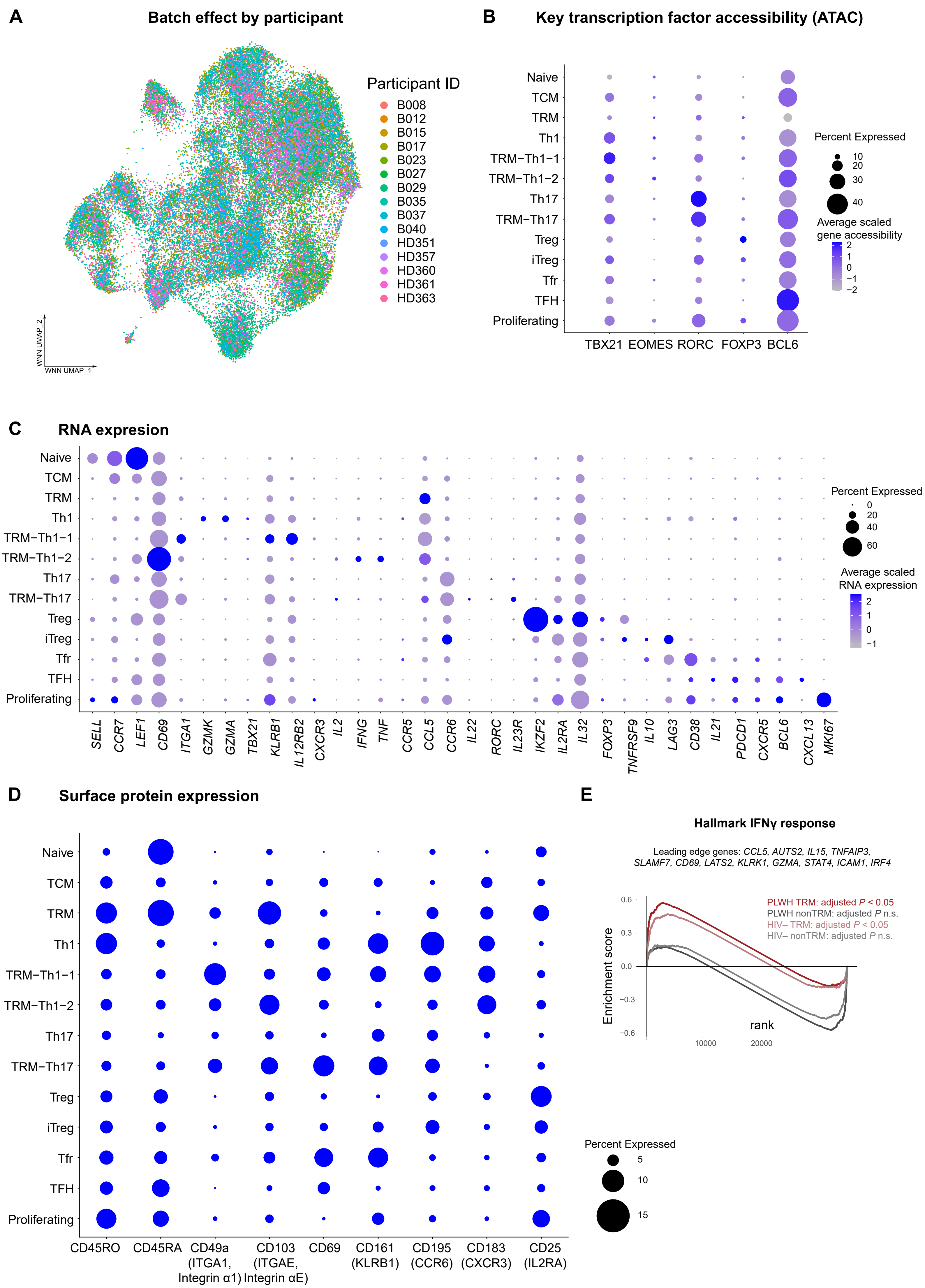

A
Batch effect by participant
Participant ID
B008
B012
B015
B017
B023
B027
B029
B035
B037
B040
HD351
HD357
HD360
HD361
HD363
WNN UMAP_2
WNN UMAP_1
B
Key transcription factor accessibility (ATAC)
Naive
TCM
TRM
Th1
TRM−Th1−1
TRM−Th1−2
Th17
TRM−Th17
Treg
iTreg
Tfr
TFH
Proliferating
TBX21 EOMES RORC FOXP3 BCL6
Percent Expressed
10
20
30
40
Average scaled gene accessibility
2
1
0
−1
−2
C
RNA expresion
SELL CCR7 LEF1 CD69 ITGA1 GZMK GZMA TBX21 KLRB1 IL12RB2 CXCR3 IL2 IFNG TNF CCR5 CCL5 CCR6 IL22 RORC IL23R IKZF2 IL2RA IL32 FOXP3 TNFRSF9 IL10 LAG3 CD38 IL21 PDCD1 CXCR5 BCL6 CXCL13 MKI67
Percent Expressed
0
20
40
60
Average scaled RNA expression
2
1
0
−1
D
Surface protein expression
CD45RO
CD45RA
CD49a (ITGA1, Integrin α1)
CD103 (ITGAE, Integrin αE)
CD69
CD161 (KLRB1)
CD195 (CCR6)
CD183 (CXCR3)
CD25 (IL2RA)
Percent Expressed
5
10
15
E
Hallmark IFNγ response
Leading edge genes: CCL5, AUTS2, IL15, TNFAIP3, SLAMF7, CD69, LATS2, KLRK1, GZMA, STAT4, ICAM1, IRF4
PLWH TRM: adjusted P < 0.05
PLWH nonTRM: adjusted P n.s.
HIV– TRM: adjusted P < 0.05
HIV– nonTRM: adjusted P n.s.
Enrichment score
0.6
0.3
0.0
−0.3
−0.6
rank
10000
20000

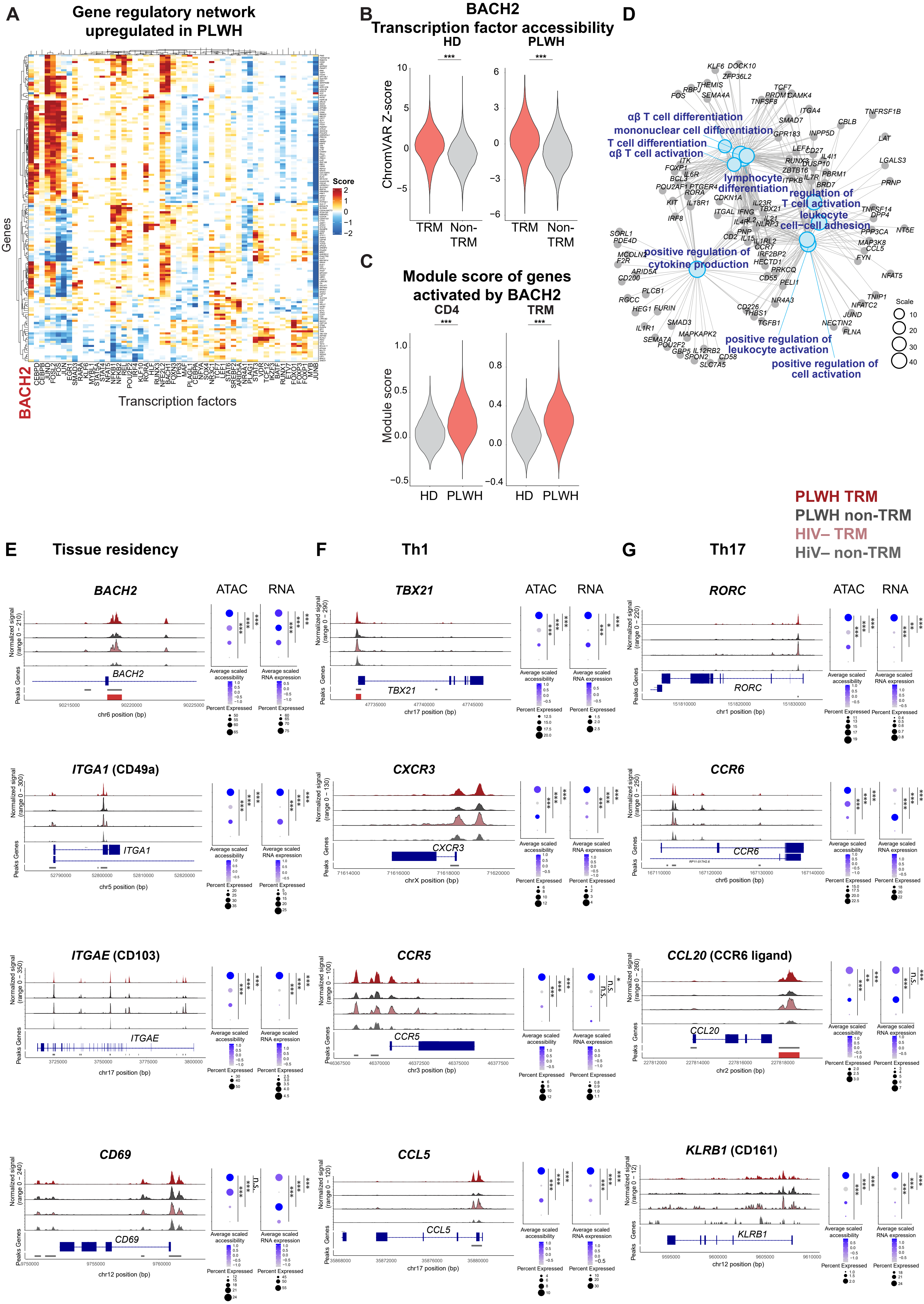

A
Gene regulatory network upregulated in PLWH
Score
Genes
BACH2
Transcription factors
B
BACH2
Transcription factor accessibility
HD
PLWH
ChromVAR Z-score
TRM
Non-TRM
C
Module score of genes activated by BACH2
CD4
TRM
Module score
HD
PLWH
D
αβ T cell differentiation
mononuclear cell differentiation
T cell differentiation
αβ T cell activation
lymphocyte differentiation
regulation of T cell activation
leukocyte cell−cell adhesion
positive regulation of cytokine production
positive regulation of leukocyte activation
positive regulation of cell activation
Scale
PLWH TRM
PLWH non-TRM
HIV– TRM
HiV– non-TRM
E
Tissue residency
BACH2
ITGA1 (CD49a)
ITGAE (CD103)
CD69
F
Th1
TBX21
CXCR3
CCR5
CCL5
G
Th17
RORC
CCR6
CCL20 (CCR6 ligand)
KLRB1 (CD161)
ATAC
RNA
Normalized signal
Peaks
Genes
Average scaled accessibility
Average scaled RNA expression
Percent Expressed

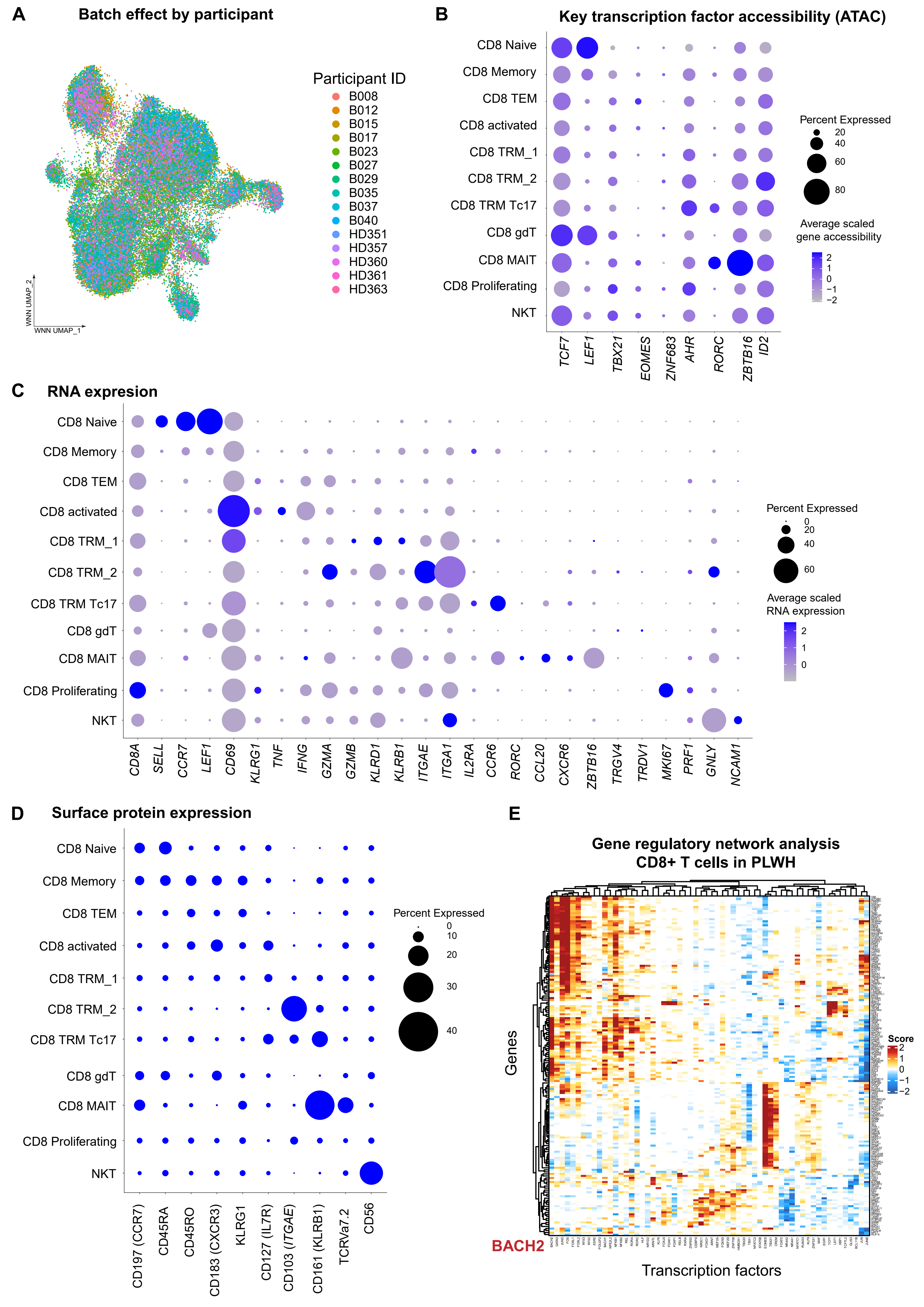
A
Batch effect by participant
Participant ID
B008
B012
B015
B017
B023
B027
B029
B035
B037
B040
HD351
HD357
HD360
HD361
HD363
WNN UMAP_2
WNN UMAP_1
B
Key transcription factor accessibility (ATAC)
CD8 Naive
CD8 Memory
CD8 TEM
CD8 activated
CD8 TRM_1
CD8 TRM_2
CD8 TRM Tc17
CD8 gdT
CD8 MAIT
CD8 Proliferating
NKT
TCF7
LEF1
TBX21
EOMES
ZNF683
AHR
RORC
ZBTB16
ID2
Percent Expressed
20
40
60
80
Average scaled gene accessibility
2
1
0
−1
−2
C
RNA expression
CD8 Naive
CD8 Memory
CD8 TEM
CD8 activated
CD8 TRM_1
CD8 TRM_2
CD8 TRM Tc17
CD8 gdT
CD8 MAIT
CD8 Proliferating
NKT
CD8A
SELL
CCR7
LEF1
CD69
KLRG1
TNF
IFNG
GZMA
GZMB
KLRD1
KLRB1
ITGAE
ITGA1
IL2RA
CCR6
RORC
CCL20
CXCR6
ZBTB16
TRGV4
TRDV1
MKI67
PRF1
GNLY
NCAM1
Percent Expressed
0
20
40
60
Average scaled RNA expression
2
1
0
D
Surface protein expression
CD8 Naive
CD8 Memory
CD8 TEM
CD8 activated
CD8 TRM_1
CD8 TRM_2
CD8 TRM Tc17
CD8 gdT
CD8 MAIT
CD8 Proliferating
NKT
CD197 (CCR7)
CD45RA
CD45RO
CD183 (CXCR3)
KLRG1
CD127 (IL7R)
CD103 (ITGAE)
CD161 (KLRB1)
TCRVa7.2
CD56
Percent Expressed
0
10
20
30
40
E
Gene regulatory network analysis
CD8+ T cells in PLWH
Genes
Score
2
1
0
−1
−2
BACH2
Transcription factors

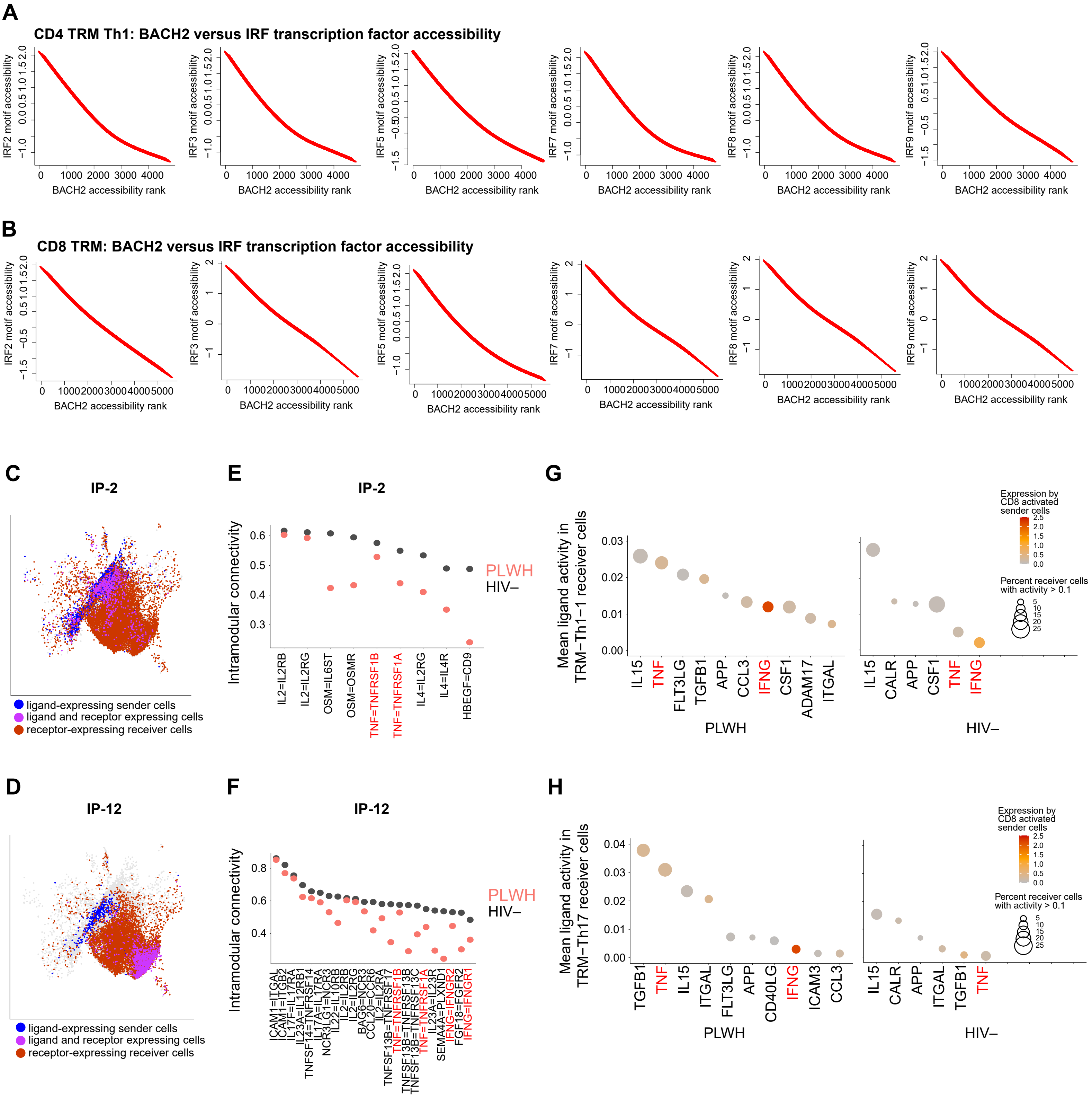

A
CD4 TRM Th1: BACH2 versus IRF transcription factor accessibility
IRF2 motif accessibility
IRF3 motif accessibility
IRF5 motif accessibility
IRF7 motif accessibility
IRF8 motif accessibility
IRF9 motif accessibility
BACH2 accessibility rank
B
CD8 TRM: BACH2 versus IRF transcription factor accessibility
C
IP-2
ligand-expressing sender cells
ligand and receptor expressing cells
receptor-expressing receiver cells
D
IP-12
E
IP-2
Intramodular connectivity
PLWH
HIV–
IL2=IL2RB
IL2=IL2RG
OSM=IL6ST
OSM=OSMR
TNF=TNFRSF1B
TNF=TNFRSF1A
IL4=IL2RG
IL4=IL4R
HBEGF=CD9
F
IP-12
G
Mean ligand activity in TRM−Th1−1 receiver cells
IL15
TNF
FLT3LG
TGFB1
APP
CCL3
IFNG
CSF1
ADAM17
ITGAL
CALR
Expression by CD8 activated sender cells
Percent receiver cells with activity > 0.1
H
Mean ligand activity in TRM−Th17 receiver cells
CD40LG
ICAM3

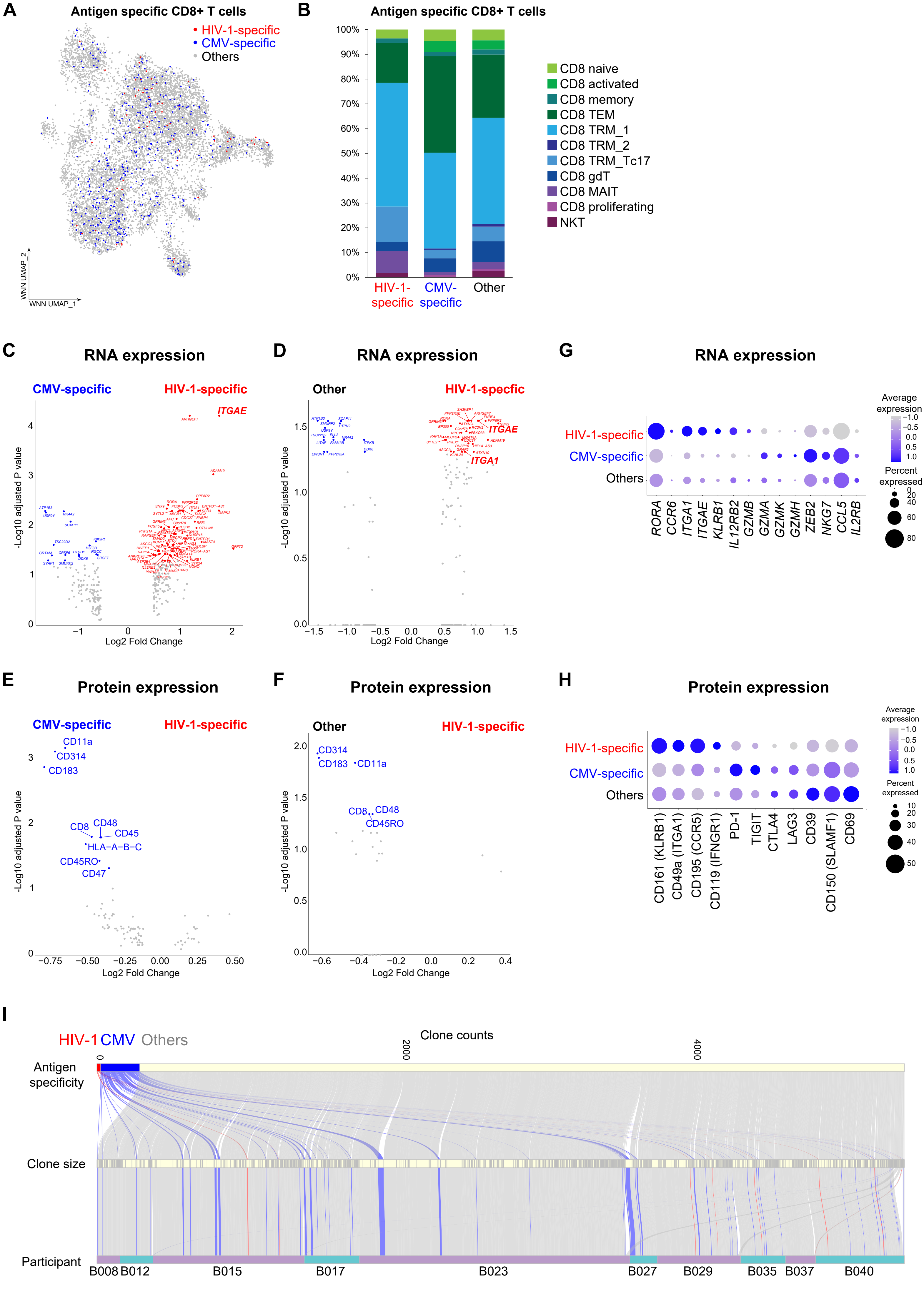
A
Antigen specific CD8+ T cells
HIV-1-specific
CMV-specific
Others
WNN UMAP_2
WNN UMAP_1
B
Antigen specific CD8+ T cells
100%
90%
80%
70%
60%
50%
40%
30%
20%
10%
0%
HIV-1-specific
CMV-specific
Other
CD8 naive
CD8 activated
CD8 memory
CD8 TEM
CD8 TRM_1
CD8 TRM_2
CD8 TRM_Tc17
CD8 gdT
CD8 MAIT
CD8 proliferating
NKT
C
RNA expression
CMV-specific
HIV-1-specific
ITGAE
-Log10 adjusted P value
Log2 Fold Change
D
RNA expression
Other
HIV-1-specfic
ITGAE
ITGA1
-Log10 adjusted P value
Log2 Fold Change
G
RNA expression
HIV-1-specific
CMV-specific
Others
RORA
CCR6
ITGA1
ITGAE
KLRB1
IL12RB2
GZMB
GZMA
GZMK
GZMH
ZEB2
NKG7
CCL5
IL2RB
Average expression
Percent expressed
E
Protein expression
CMV-specific
HIV-1-specific
CD11a
CD314
CD183
CD8
CD48
CD45
HLA-A-B-C
CD45RO
CD47
-Log10 adjusted P value
Log2 Fold Change
F
Protein expression
Other
HIV-1-specific
CD314
CD183
CD11a
CD8
CD48
CD45RO
-Log10 adjusted P value
Log2 Fold Change
H
Protein expression
HIV-1-specific
CMV-specific
Others
CD161 (KLRB1)
CD49a (ITGA1)
CD195 (CCR5)
CD119 (IFNGR1)
PD-1
TIGIT
CTLA4
LAG3
CD39
CD150 (SLAMF1)
CD69
Average expression
Percent expressed
I
HIV-1 CMV Others
Clone counts
0
2000
4000
Antigen specificity
Clone size
Participant
B008
B012
B015
B017
B023
B027
B029
B035
B037
B040

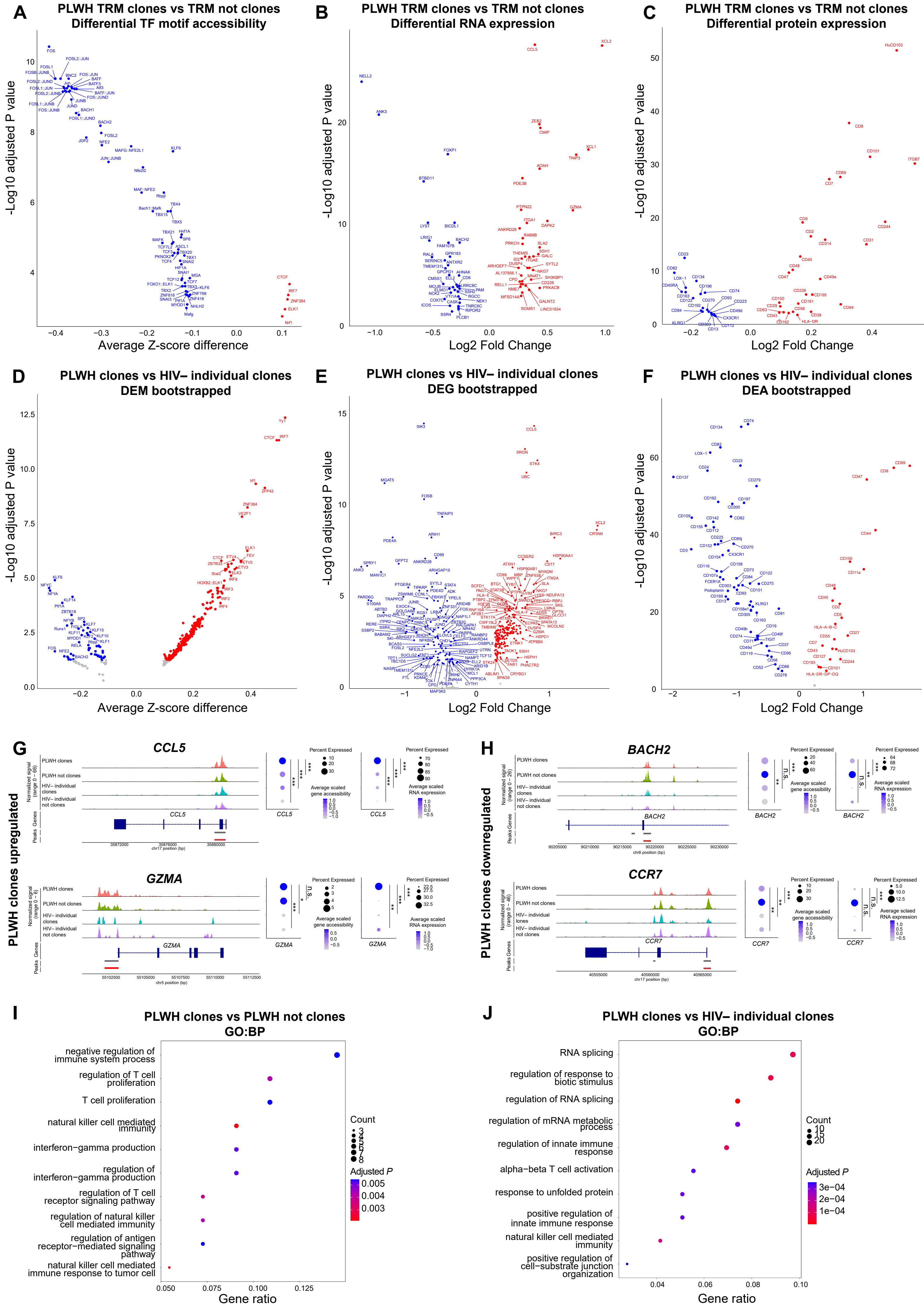
A
PLWH TRM clones vs TRM not clones
Differential TF motif accessibility
-Log10 adjusted P value
Average Z-score difference
B
PLWH TRM clones vs TRM not clones
Differential RNA expression
-Log10 adjusted P value
Log2 Fold Change
C
PLWH TRM clones vs TRM not clones
Differential protein expression
-Log10 adjusted P value
Log2 Fold Change
D
PLWH clones vs HIV− individual clones
DEM bootstrapped
-Log10 adjusted P value
Average Z-score difference
E
PLWH clones vs HIV− individual clones
DEG bootstrapped
-Log10 adjusted P value
Log2 Fold Change
F
PLWH clones vs HIV− individual clones
DEA bootstrapped
-Log10 adjusted P value
Log2 Fold Change
G
PLWH clones upregulated
CCL5
GZMA
PLWH clones
PLWH not clones
HIV− individual clones
HIV− individual not clones
Normalized signal
Peaks
Genes
Percent Expressed
Average scaled gene accessibility
Average scaled RNA expression
H
PLWH clones downregulated
BACH2
CCR7
I
PLWH clones vs PLWH not clones
GO:BP
negative regulation of immune system process
regulation of T cell proliferation
T cell proliferation
natural killer cell mediated immunity
interferon−gamma production
regulation of interferon−gamma production
regulation of T cell receptor signaling pathway
regulation of natural killer cell mediated immunity
regulation of antigen receptor−mediated signaling pathway
natural killer cell mediated immune response to tumor cell
Gene ratio
Count
Adjusted P
J
PLWH clones vs HIV− individual clones
GO:BP
RNA splicing
regulation of response to biotic stimulus
regulation of RNA splicing
regulation of mRNA metabolic process
regulation of innate immune response
alpha−beta T cell activation
response to unfolded protein
positive regulation of innate immune response
natural killer cell mediated immunity
positive regulation of cell−substrate junction organization
Gene ratio
Count
Adjusted P

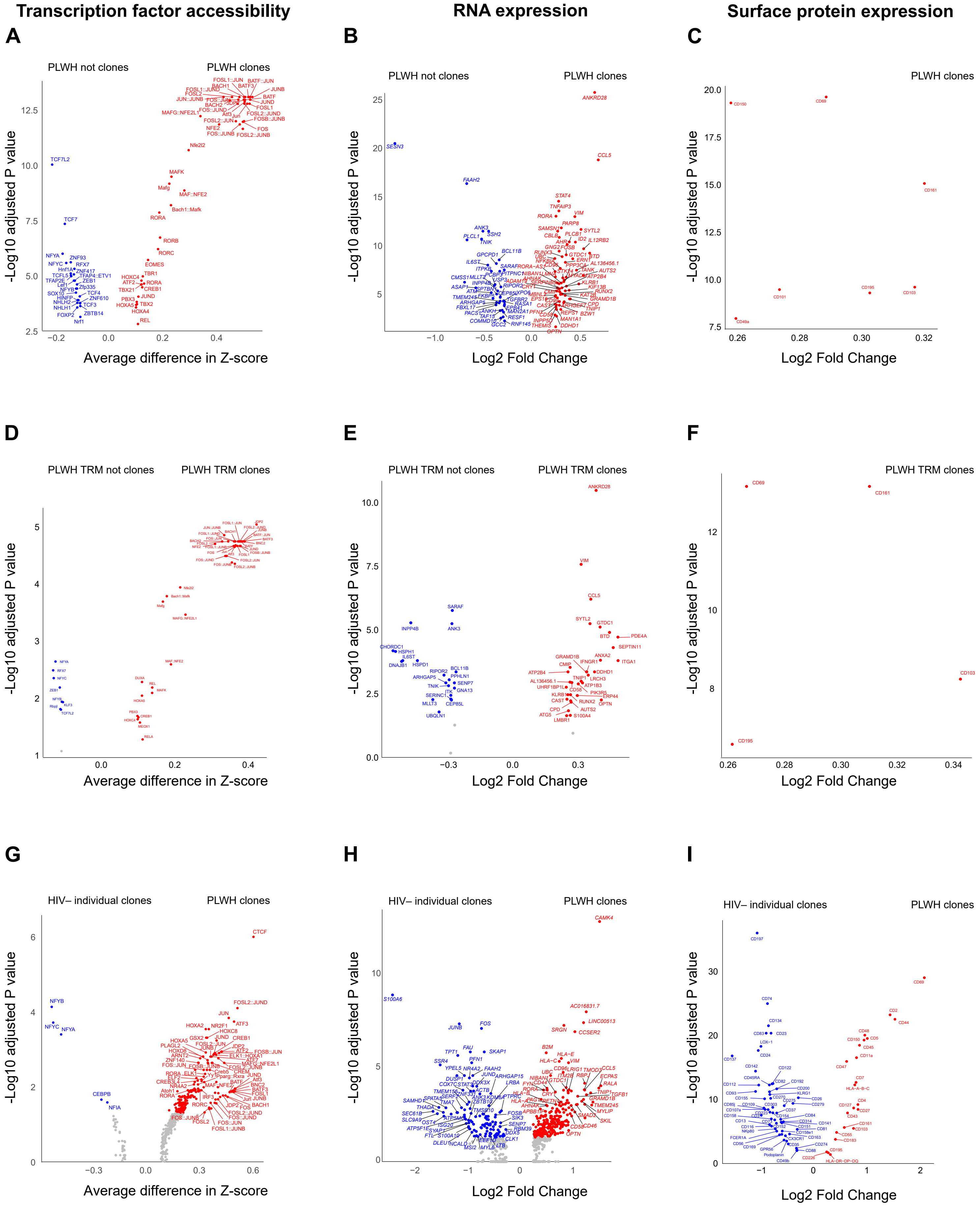

Transcription factor accessibility
RNA expression
Surface protein expression
A
B
C
PLWH not clones
PLWH clones
-Log10 adjusted P value
Average difference in Z-score
Log2 Fold Change
D
E
F
PLWH TRM not clones
PLWH TRM clones
G
H
I
HIV– individual clones
PLWH clones

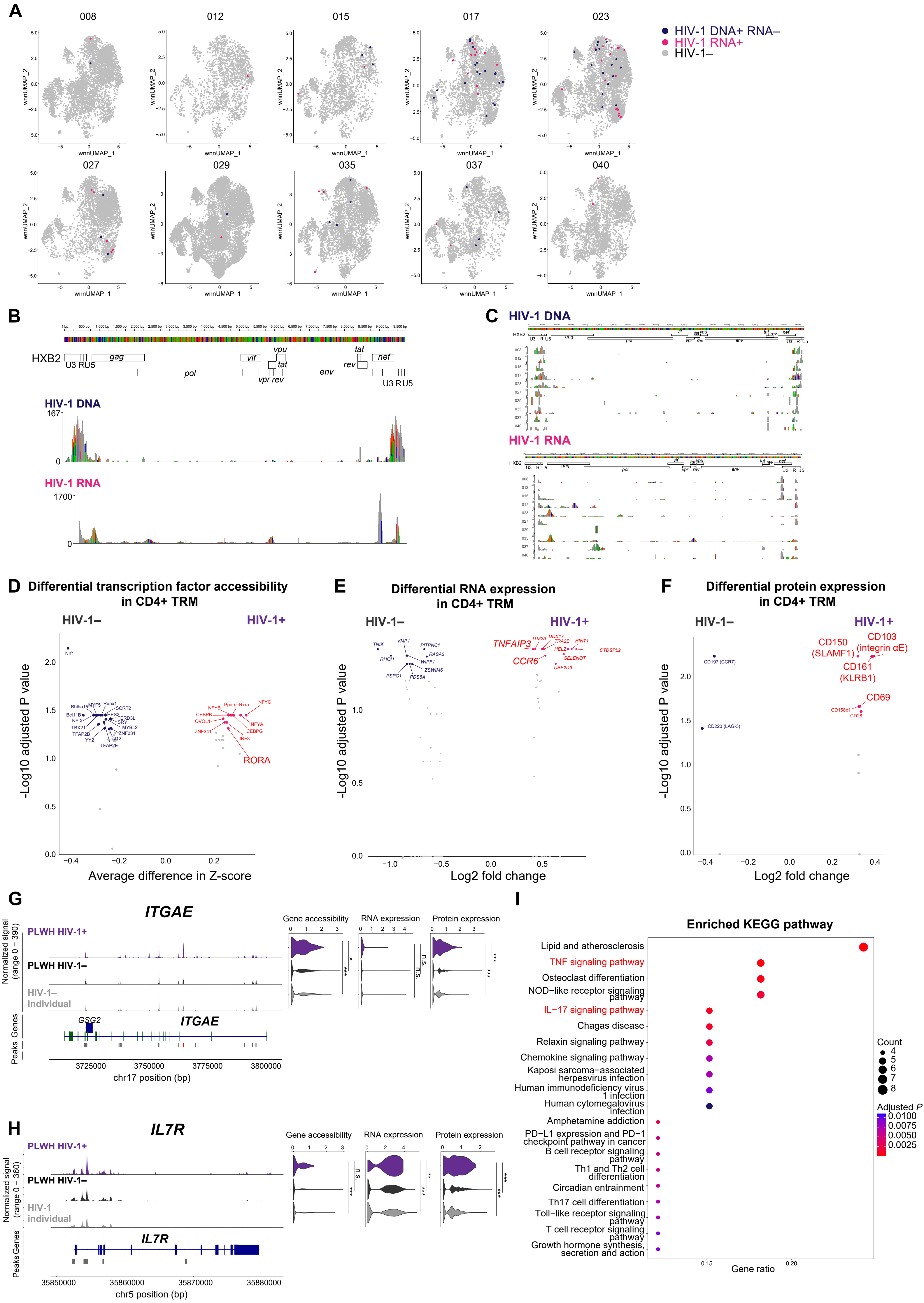
A
008
012
015
017
023
027
029
035
037
040
wnnUMAP_1
wnnUMAP_2
HIV-1 DNA+ RNA–
HIV-1 RNA+
HIV-1–
B
HXB2
U3 RU5
gag
pol
vif
vpu
tat
vpr rev
rev
env
tat
nef
U3 R U5
HIV-1 DNA
167
0
HIV-1 RNA
1700
0
C
HIV-1 DNA
HIV-1 RNA
D
Differential transcription factor accessibility in CD4+ TRM
HIV-1–
HIV-1+
Nrf1
RORA
-Log10 adjusted P value
Average difference in Z-score
E
Differential RNA expression in CD4+ TRM
HIV-1–
HIV-1+
TNFAIP3
CCR6
-Log10 adjusted P value
Log2 fold change
F
Differential protein expression in CD4+ TRM
HIV-1–
HIV-1+
CD197 (CCR7)
CD223 (LAG-3)
CD150 (SLAMF1)
CD103 (integrin αE)
CD161 (KLRB1)
CD69
CD158e1
CD28
-Log10 adjusted P value
Log2 fold change
G
ITGAE
PLWH HIV-1+
PLWH HIV-1–
HIV-1– individual
Normalized signal (range 0 – 390)
Gene accessibility
RNA expression
Protein expression
GSG2
ITGAE
Genes
Peaks
3725000
3750000
3775000
3800000
chr17 position (bp)
H
IL7R
PLWH HIV-1+
PLWH HIV-1–
HIV-1 individual
Normalized signal (range 0 – 360)
Gene accessibility
RNA expression
Protein expression
IL7R
Genes
Peaks
35850000
35860000
35870000
35880000
chr5 position (bp)
I
Enriched KEGG pathway
Lipid and atherosclerosis
TNF signaling pathway
Osteoclast differentiation
NOD−like receptor signaling pathway
IL−17 signaling pathway
Chagas disease
Relaxin signaling pathway
Chemokine signaling pathway
Kaposi sarcoma−associated herpesvirus infection
Human immunodeficiency virus 1 infection
Human cytomegalovirus infection
Amphetamine addiction
PD−L1 expression and PD−1 checkpoint pathway in cancer
B cell receptor signaling pathway
Th1 and Th2 cell differentiation
Circadian entrainment
Th17 cell differentiation
Toll−like receptor signaling pathway
T cell receptor signaling pathway
Growth hormone synthesis, secretion and action
Count
Adjusted P
Gene ratio

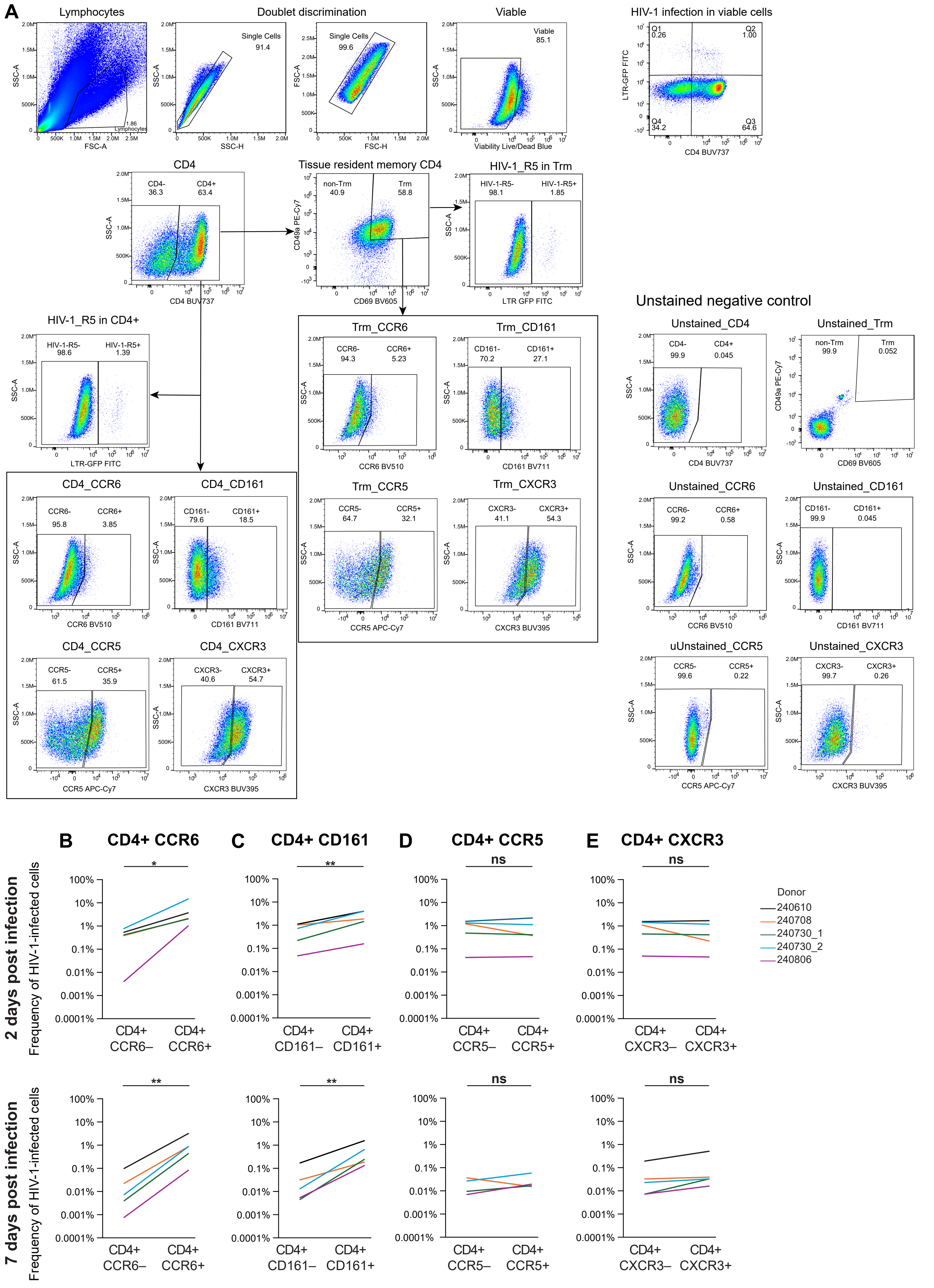
A
Lymphocytes
Doublet discrimination
Viable
HIV-1 infection in viable cells
Single Cells 91.4
Single Cells 99.6
Viable 85.1
1.86 Lymphocytes
FSC-A
SSC-A
SSC-H
FSC-H
Viability Live/Dead Blue
LTR-GFP FITC
CD4 BUV737
Q1 0.26
Q2 1.00
Q3 64.6
Q4 34.2
CD4
CD4- 36.3
CD4+ 63.4
Tissue resident memory CD4
non-Trm 40.9
Trm 58.8
CD49a PE-Cy7
CD69 BV605
HIV-1_R5 in Trm
HIV-1-R5- 98.1
HIV-1-R5+ 1.85
LTR GFP FITC
HIV-1_R5 in CD4+
HIV-1-R5- 98.6
HIV-1-R5+ 1.39
LTR-GFP FITC
Trm_CCR6
CCR6- 94.3
CCR6+ 5.23
CCR6 BV510
Trm_CD161
CD161- 70.2
CD161+ 27.1
CD161 BV711
Trm_CCR5
CCR5- 64.7
CCR5+ 32.1
CCR5 APC-Cy7
Trm_CXCR3
CXCR3- 41.1
CXCR3+ 54.3
CXCR3 BUV395
CD4_CCR6
CCR6- 95.8
CCR6+ 3.85
CD4_CD161
CD161- 79.6
CD161+ 18.5
CD4_CCR5
CCR5- 61.5
CCR5+ 35.9
CD4_CXCR3
CXCR3- 40.6
CXCR3+ 54.7
Unstained negative control
Unstained_CD4
CD4- 99.9
CD4+ 0.045
Unstained_Trm
non-Trm 99.9
Trm 0.052
Unstained_CCR6
CCR6- 99.2
CCR6+ 0.58
Unstained_CD161
CD161- 99.9
CD161+ 0.045
uUnstained_CCR5
CCR5- 99.6
CCR5+ 0.22
Unstained_CXCR3
CXCR3- 99.7
CXCR3+ 0.26
B CD4+ CCR6
C CD4+ CD161
D CD4+ CCR5
E CD4+ CXCR3
*
**
ns
2 days post infection
7 days post infection
Frequency of HIV-1-infected cells
100%
10%
1%
0.1%
0.01%
0.001%
0.0001%
CD4+ CCR6–
CD4+ CCR6+
CD4+ CD161–
CD4+ CD161+
CD4+ CCR5–
CD4+ CCR5+
CD4+ CXCR3–
CD4+ CXCR3+
Donor
240610
240708
240730_1
240730_2
240806

### Table S1. Clinical characteristics of study participants

| Participant | Age | Sex | Race | Ethnicity | ART | Duration of ART (months) | CD4 count (cells/µl) | HIV viral load (copies/ml) |
|---|---|---|---|---|---|---|---|---|
| People living with HIV (PLWH) | | | | | | | | |
| 008 | 48 | Male | White | Not Hispanic | FTC/RPV/TDF | 48.8 | 1247 | <20 |
| 012 | 47 | Female | Black | Not Hispanic | EVG/c/FTC/FTC/TDF | 26.3 | 731 | <20 |
| 015 | 58 | Male | White | Not Hispanic | DOL/ABC/3TC | 8.6 | 457 | <20 |
| 017 | 49 | Male | Black | Not Hispanic | FTC/RPV/TDF | 20.9 | 531 | <20 |
| 023 | 45 | Male | Black | Not Hispanic | FTC/PRV/TDF | 50.8 | 1087 | <20 |
| 027 | 41 | Male | White | Not Hispanic | EFV/FTC/TDF | 86.8 | 584 | <20 |
| 029 | 28 | Male | Black | Not Hispanic | EFV/FTC/TDF | 50.1 | 1146 | <20 |
| 035 | 37 | Male | Other | Hispanic | EFV/FTC/TDF | 74.4 | 732 | <20 |
| 037 | 32 | Male | Black | Not Hispanic | FTC/RPV/TDF | 25.0 | 884 | <20 |
| 040 | 41 | Male | White | Not Hispanic | FTC/RPV/TAF | 2.4 | 538 | 178 |
| People living without HIV (HD) | | | | | | | | |
| 351 | 37 | Male | White | Hispanic | | | | |
| 357 | 51 | Male | Black | Hispanic | | | | |
| 360 | 38 | Male | Asian | Hispanic | | | | |
| 361 | 27 | Female | White | Hispanic | | | | |
| 363 | 25 | Female | White | Hispanic | | | | |

Surgical resection from people living without HIV for *in vitro* validation

| Participant | Age | Sex | Location | Diagnosis |
|---|---|---|---|---|
| 240610 | 57 | Male | Right normal colon | colon cancer |
| 240708 | 74 | Female | Left normal colon | diverticulitis |
| 240730 | 81 | Female | Right normal colon | colon cancer |
| 240806 | 64 | Female | Right normal colon | colon cancer |

3TC, lamivudine; ABC, abacavir; /c, cobicistat; DOL, dolutegravir; EFV, efavirenz; EVG, elvitegravir; FTC, emtricitabine; RPV, rilpivirine; TAF, tenofovir alafenamide; TDF, tenofovir
